# Chameleonicity Beyond Compactness: Conformational Persistence and Passive Permeability Across 20 PROTAC Ensembles

**DOI:** 10.64898/2026.10.05.756765

**Authors:** Farzaneh Jalalypour, Rocío Mercado

## Abstract

Proteolysis-targeting chimeras (PROTACs) are large, flexible molecules that frequently extend beyond conventional drug-like chemical space, including Lipinski’s rule-of-five boundaries. Their membrane permeability has often been linked to molecular chameleonicity and the ability to adopt compact conformations with reduced polar surface exposure; however, the underlying conformational determinants remain unclear. Here, we investigated 20 PROTACs spanning distinct passive-permeability classes using enhanced-sampling molecular dynamics (ES-MD). All permeability classes accessed compact states with low radius of gyration (Rg) and polar surface area (PSA), indicating that compact-state accessibility alone does not distinguish permeable from non-permeable PROTACs. We therefore introduce maximum ring size (MRS), defined as the size of the largest intramolecular hydrogen-bonded ring, as a metric for identifying compact conformations stabilized by long-range intramolecular hydrogen bonding. Like the other descriptors examined, MRS neither separates nor ranks the permeability classes, but helps locate folded states in the large ensemble. Representative compact conformations from selected highly permeable, moderately permeable, and non-permeable compounds were subsequently examined by all-atom MD in water and chloroform. The highly permeable PROTAC held a closed-flat ensemble in both environments, but in two structurally distinct arrangements, whereas the other two redistributed across heterogeneous ensembles. Passive diffusion therefore appears to depend on the population, stability, and environment-dependent persistence of permeable-like states rather than on compactness alone.

## 1 Introduction

For decades, drug discovery has relied on the identification of active or allosteric protein sites where ligands can bind and modulate the function of disease-related proteins. Although highly successful, this approach is limited to proteins with tractable binding pockets, leaving many clinically relevant targets beyond the reach of conventional strategies. Proteolysis-targeting chimeras (PROTACs) offer an alternative modality by acting as degradation-inducing molecular tags that recruit both a protein of interest and an E3 ubiquitin ligase, leading to ubiquitination and proteasomal elimination of the target. By inducing selective protein degradation rather than directly modulating protein activity through a conventional binding site, PROTACs expand therapeutic access to targets that have traditionally been considered “undruggable” [1, 2]. They have also shown success in achieving rapid and reversible protein knockdown as an alternative to CRISPR/Cas-based approach [3]. However, the intracellular efficacy of PROTACs depends strongly on their ability to cross cellular membranes and reach the sites-of-action, making permeability a major challenge in degrader design. Owing to their heterobifunctional architecture, PROTACs typically exhibit high molecular weight, considerable polarity, multiple hydrogen-bonding groups, and substantial conformational flexibility. Consequently, they often fall outside the physicochemical boundaries defined by Lipinski’s rule of five [4] and Veber’s rules [5], which are commonly used as traditional guidelines for drug-likeness. Instead, PROTACs occupy beyond-rule-of-five (bRo5) chemical space, where physicochemical descriptors often exceed the ranges typically associated with orally available small-molecule drugs. Nevertheless, the existence of orally bioavailable compounds within bRo5 chemical space indicates that, although PROTAC design remains challenging, it should not be constrained solely by classical physicochemical rules [6, 7]. Molecular chameleonicity has been proposed as one possible explanation for the passive permeability of beyond-rule-of-five (bRo5) molecules, including PROTACs [8, 9]. This concept refers to the ability of a molecule to alter its conformational state in response to its environment, a property first quantified for macrocycles and other high molecular weight (MW) drugs [9]. For PROTACs, such adaptation may be driven by transient intramolecular interactions that shield polar groups and reduce solvent-exposed polar surface area, a behavior demonstrated from solution conformational ensembles by Atilaw et al. [10]. The resulting compact, polarity-shielded conformations, often reflected by a reduced radius of gyration (Rg) and 3D polar surface area (PSA), may enhance passive diffusion across cellular membranes and thereby support cellular uptake [11, 12]. However, the mechanistic relationship between chameleonic behavior and PROTAC permeability remains poorly understood.

Chameleonicity can be quantified using several complementary approaches. Structural techniques such as X-ray crystallography and NMR spectroscopy can be used to characterize conformational ensembles, which can subsequently be analyzed through conformer-specific molecular descriptors [10, 13]. In parallel, chromatography-based descriptors offer a high-throughput strategy for assessing environment-dependent molecular behavior. Among these, ChamelogD and Chamelogk have been introduced as chromatographic metrics for evaluating molecular chameleonicity [14]. More recently, Kim et al. [15] combined conventional 2D descriptors with 3D descriptors derived from well-tempered metadynamics ensembles, including Rg, 3D PSA, and the number of intramolecular hydrogen bonds, and reported improved machine-learning prediction of heterobifunctional degrader permeability. In that work, as in the conformational studies above, intramolecular hydrogen bonding enters only as a count: how far apart the donor and acceptor sit along the covalent path, and therefore how much of the structure a single interaction closes into a ring, is not captured by any descriptor used to date.

At present, no single molecular descriptor can reliably explain or predict passive permeability, or fully capture its relationship with chameleonicity, highlighting the need to consider multiple complementary physicochemical and conformational metrics rather than relying on one parameter alone. Accordingly, several studies have sought to develop multiparameter guidelines and scoring systems. Examples include AbbVie’s AB-MPS score; which integrates lipophilicity, aromaticity, and molecular flexibility [16]; EPSA-to-TPSA ratio (ETR), the new experimental “chameleonicity” parameter and polarity reduction descriptor [17]; the bifunctional bioavailability index (BBI) [18]; and the balanced permeability index (BPI) [19]. Together, these approaches aim to capture the polarity exposure, chameleonicity, conformational behavior, and balance between size, lipophilicity, molecular shape, and permeability-relevant properties. However, none of these single descriptors was sufficient to rationalize PROTAC permeability or distinguish permeable from poorly permeable ones [20].

A detailed understanding of PROTAC dynamics, together with characterization of their solution ensembles, may therefore provide important insights into their permeability behavior and help refine spatial descriptors and bioavailability-related metrics. Several experimental and computational studies have investigated PROTACs as integral components of ternary complexes, where they function as dynamic molecular bridges between the target protein and the recruited E3 ligase. These studies primarily focus on the bound conformation of the PROTAC within the protein complex and examine how linker geometry, degrader flexibility, protein–protein contacts, and ternary-complex stability influence productive complex formation and degradation efficiency.[21–25] Complementary to this perspective, the analysis of isolated PROTACs in solution provides important insight into permeability-relevant conformational behavior before target engagement. For example, Abeje et al. [11] investigated a series of nine VHL-based PROTACs with systematic variations in linker structure to determine how linker composition influences passive cell permeability. The study showed that relatively small changes in linker design can markedly affect permeability by altering the conformational preferences of the degrader in solution. Highly permeable PROTACs were associated with linkers that promoted folded conformations and reduced solvent-exposed polarity, while all compounds exhibited low aqueous solubility regardless of their permeability.

In this study, we investigated the relationship between PROTAC conformational behavior and passive permeability using a curated dataset of 20 PROTACs with reported parallel artificial membrane permeability assay (PAMPA) values, classified into permeable (P), moderately permeable (M), and non-permeable (NP) groups. Conformational ensembles were generated using enhanced sampling methods for molecular dynamics (ES-MD) with Rg used as a collective variable (Moltiverse) to promote sampling of compact and extended states. The resulting ensembles were analyzed using multiple structural and physicochemical descriptors to identify potential relations. Although all PROTACs were able to access compact conformations in all groups, no single descriptor showed a strong overall correlation or pattern with permeability. We introduce the maximum intramolecular hydrogen-bond ring-size (MRS) descriptor to identify compact conformers. MRS is the size of the largest pseudo-ring formed by an intramolecular hydrogen bond, counting the bridging hydrogen, donor, acceptor, and atoms along the connecting covalent path. Unlike IMHB counts, MRS distinguishes local hydrogen bonds from those that bring distant molecular regions together. MRS did not discriminate between permeability classes in our analysis; however, it provided a criterion for selecting comparable compact structures across classes. For three PROTACs representing different permeability classes, PROTAC-1 (P), PROTAC-3 (M), and PROTAC-18 (NP), conformers with greater MRS were selected from the ES-MD-generated ensembles as putative folded, polarity-shielded states. Their stability and solvent-dependent conformational adaptation were subsequently evaluated using unbiased molecular dynamics (U-MD) simulations in polar and apolar environments, providing an indirect assessment of chameleonic behavior. Our results suggest that no single conformational feature directly determines PROTAC permeability. Molecular chameleonicity should therefore not be viewed simply as the ability to adopt a compact, permeable state; rather, permeability may depend on the population, stability, and environment-dependent persistence of specific permeable-like conformations. We further introduce maximum ring size as a practical metric for selecting compact structures stabilized by long-range intramolecular hydrogen bonding. In addition, enhanced-sampling MD using Rg as the collective variable provides a practical framework for generating diverse PROTAC conformational ensembles and supporting future studies of rational degrader design.

Our main contributions are:

- We provide conformational ensembles for 20 PROTACs spanning different permeability classes and demonstrate the utility of Moltiverse for sampling PROTACs diverse conformational states.
- We introduce maximum intramolecular hydrogen-bond ring-size (MRS) as an interaction-based criterion for locating compact conformers stabilized by long-range intramolecular hydrogen bonding. MRS does not correlate with permeability; its value is in selecting comparably folded starting structures from large ensembles.
- We show that PROTAC permeability is not explained by conventional molecular descriptors, and that compact permeable-like conformations are accessible to all permeability classes.
- We show that the permeable PROTAC occupies a closed-flat ensemble in both polar and apolar media, yet in two structurally distinct arrangements, indicating that environmental adaptation need not proceed through compaction–extension cycling, as classically assumed.

## 2 Methods

### 2.1 PROTAC dataset

Given the limited availability of experimentally reported PAMPA data for PROTAC molecules, a curated dataset comprising 20 PROTAC compounds was compiled from the literature. PAMPA is a high-throughput *in vitro* assay that measures passive permeability across a lipid-infused artificial membrane, independent of proteins, active transport, efflux and metabolism.

Different PAMPA models are used, in which the artificial membrane composition, buffer system, and incubation conditions are modified to mimic specific biological barriers, such as the gastrointestinal tract or the blood–brain barrier. The BBB-PAMPA model, typically based on porcine polar brain lipids, represents a more restrictive barrier than the GIT-PAMPA model. Although these models may show qualitative agreement, their results do not necessarily correlate linearly or preserve the same rank order. Compounds with measurable BBB-PAMPA permeability may exhibit detectable GIT-PAMPA permeability, but this should be interpreted as a general trend rather than an absolute rule [26, 27].

The corresponding PAMPA permeability values and SMILES were obtained from PROTAC-DB [28], with the exception of PROTAC-3, for which the chemical structure reported by Atilaw et al. [10] was manually converted into a SMILES representation. The PAMPA permeability data were curated and converted to consistent units; therefore, the values reported here may differ from those originally listed in PROTAC-DB. The SMILES strings were retained as provided by PROTAC-DB and used as the initial molecular representations for all subsequent computational analyses, including descriptor calculation, and enhanced sampling simulations. For each PROTAC, we computed 2D molecular descriptors using RDKit 2025 [29]. Hydrogen atoms were retained and explicitly included in all calculations. We provide the full descriptor dataset in Table 1, while the full supplementary dataset is available through Zenodo.

**Table 1.** Complete descriptor dataset. P: permeable; M: moderately permeable; NP: non-permeable. Hydrogens included. PROTAC-1 and PROTAC-2 were measured using BBB-PAMPA assays. Detailed assay-type annotations for each compound are provided in the supplementary data.

| PROTAC ID (Class) | MW | HAC | Rings | HBA | HBD | RB | TPSA(Å<sup>2</sup>) | cLogP | PAMPA (nm/s) | Source |
| --- | --- | --- | --- | --- | --- | --- | --- | --- | --- | --- |
| 1 (P) | 801.64 | 56 | 7 | 11 | 4 | 15 | 194.24 | 3.49 | 206.8 | [30] |
| 2 (P) | 857.75 | 60 | 7 | 11 | 4 | 19 | 194.24 | 4.92 | 153.3 | [30] |
| 3 (M) | 1034.13 | 73 | 7 | 15 | 4 | 20 | 219.56 | 6.35 | 14.1 | [10] |
| 4 (M) | 973.62 | 67 | 7 | 15 | 3 | 18 | 199.46 | 6.04 | 6 | [31] |
| 5 (M) | 1000.69 | 69 | 7 | 14 | 4 | 20 | 202.26 | 6.18 | 5 | [31] |
| 6 (M) | 987.65 | 68 | 7 | 15 | 3 | 18 | 199.46 | 6.45 | 3 | [31] |
| 7 (M) | 986.66 | 68 | 7 | 14 | 4 | 18 | 202.26 | 5.87 | 2 | [31] |
| 8 (M) | 1001.67 | 69 | 7 | 15 | 3 | 20 | 199.46 | 6.76 | 2 | [31] |
| 9 (M) | 730.83 | 54 | 7 | 10 | 3 | 10 | 166.17 | 3.05 | 2 | [32] |
| 10 (NP) | 1003.64 | 69 | 7 | 16 | 3 | 20 | 208.69 | 5.54 | 1 | [31] |
| 11 (NP) | 972.63 | 67 | 7 | 14 | 4 | 18 | 202.26 | 5.47 | 0.8 | [31] |
| 12 (NP) | 742.79 | 55 | 8 | 11 | 2 | 8 | 174.45 | 2.77 | 0.7 | [32] |
| 13 (NP) | 742.79 | 55 | 8 | 11 | 2 | 8 | 174.45 | 2.77 | 0.5 | [32] |
| 14 (NP) | 1002.66 | 69 | 7 | 15 | 4 | 20 | 211.49 | 4.96 | 0.1 | [31] |
| 15 (NP) | 763.86 | 56 | 8 | 10 | 2 | 6 | 167.92 | 0.99 | 0.09 | [32] |
| 16 (NP) | 760.78 | 56 | 8 | 11 | 2 | 8 | 174.45 | 2.87 | 0.07 | [32] |
| 17 (NP) | 997.20 | 71 | 8 | 15 | 3 | 19 | 189.92 | 3.84 | 0.02 | [33] |
| 18 (NP) | 1041.26 | 74 | 8 | 16 | 3 | 22 | 199.15 | 3.69 | 0.01 | [33] |
| 19 (NP) | 995.23 | 71 | 8 | 14 | 3 | 19 | 180.69 | 5.06 | 0.01 | [33] |
| 20 (NP) | 763.86 | 56 | 8 | 10 | 2 | 6 | 167.92 | 0.99 | 0.007 | [32] |

### 2.2 Enhanced sampling molecular dynamics simulation (ES-MD)

PROTAC conformer ensembles were generated using Moltiverse, an enhanced-sampling molecular dynamics-based conformer generation protocol developed by Bedoya et al. [34]. Moltiverse promotes efficient exploration of molecular conformational space by biasing physically meaningful collective variables, particularly the radius of gyration, thereby enabling systematic sampling of both compact and extended conformational states. This approach helps overcome energy barriers that can limit conformational transitions in conventional molecular dynamics simulations. For each PROTAC, the curated SMILES representation was provided as input and converted into an initial 3D molecular structure. The molecules were then prepared and parameterized using the GAFF2 force field through AmberTools 24 [35, 36], followed by energy minimization to remove unfavorable contacts and generate physically reasonable starting geometries. Enhanced conformational sampling was subsequently performed in vacuum using the molecular extended adaptive biasing force approach combined with metadynamics, as implemented in NAMD 3.0.1 [37], with the Rg used as the collective variable. Biasing along the Rg enabled exploration of both compact and extended conformational states, which is particularly relevant for highly flexible PROTAC molecules. The generated trajectories were then processed to extract representative structures through hierarchical clustering using the open-source hclust library, written in the Crystal programming language [38], thereby removing redundancy and producing a diverse conformational ensemble. The selected conformers were further refined by molecular mechanics optimization and semi-empirical quantum mechanical (QM) optimization using GFN2-xTB [39]. All input files, output data, and analysis results are available in the project’s Zenodo repository.

### 2.3 Unbiased molecular dynamics simulation (U-MD)

Unbiased molecular dynamics (U-MD) simulations were conducted using AmberTools [35], for system preparation and GROMACS 2025.1 [40] for production runs. PROTAC structures were parameterized with the General Amber Force Field (GAFF2) using the antechamber module of AmberTools 24.8 [35, 41], with partial atomic charges assigned via the AM1-BCC method [42]. All compounds were simulated in their neutral form. Missing parameters were generated via parmchk2 [35]. Systems were constructed in tleap [35] by solvating each PROTAC in explicit cubic solvent boxes to represent distinct environments. Water was used to model polar conditions, while chloroform was employed for apolar conditions, with a 5 Å solvent buffer around the solute. Appropriate solvent parameters were applied and systems were neutralized by the addition of counterions where required, resulting in two simulation systems for each PROTAC. The AMBER topology and coordinate files were converted to GROMACS format using ACPYPE [43]. Following energy minimization and equilibration under constant volume (NVT) and pressure (NPT), production simulations were performed under periodic boundary conditions using a 2 fs integration time step. The pressure and temperature were maintained at 1 bar and 300 K using the C-rescale barostat [44] and V-rescale thermostat [45], respectively. All covalent bonds involving hydrogen atoms were constrained using the LINCS algorithm [46]. Long-range electrostatic interactions were treated with the particle-mesh Ewald method [47]. For each system, four independent replicas of 200 ns were carried out, resulting in a total simulation time of 800 ns per system and enabling comparative analysis of PROTAC conformational behavior in polar and apolar environments.

### 2.4 Physicochemical and conformational analysis

All molecular descriptors and trajectory-derived properties were calculated using in-house Python workflows that integrated established cheminformatics and molecular-simulation analysis libraries, including RDKit, MDTraj, and MDAnalysis [29, 48–50] and were calculated over the cumulative 800 ns ensemble.

#### 2.4.1 2D descriptors

The 2D-physicochemical descriptors including MW, exact mass, heavy atom count (HAC), ring count, hydrogen-bond acceptors (HBA), hydrogen-bond donors (HBD), rotatable bond count (RB), topological polar surface area (TPSA), and lipophilicity-related descriptors; Crippen LogP (cLogP), were calculated from SMILES.

#### 2.4.2 3D descriptors

To further characterize conformational properties of ensembles, additional 3D and trajectory-derived descriptors including radius of gyration (Rg), solvent accessible surface area (SASA), 3D polar surface area (PSA) were calculated. PSA was calculated as the summed solvent-accessible surface area of all nitrogen and oxygen atoms together with the hydrogens bonded to them; sulfur was not included.

#### 2.4.3 Intramolecular hydrogen bonding and maximum ring size metric

Intramolecular hydrogen bonds (IHBs) were identified geometrically: any hydrogen bonded to N or O lying within 3 Å of a non-bonded N or O acceptor was counted as an IHB. Donor–H···acceptor angles were recorded but not used as a filter. For each IHB, the hydrogen-bond ring (HBR) size was taken as the number of atoms in the pseudo-ring it closes, comprising the donor, acceptor, intervening path atoms, and the bridging hydrogen. The largest HBR per conformer (MRS) was used to identify folded conformations (Section 3.3).

#### 2.4.4 Structural clustering

Cluster analysis was performed to identify representative and frequently sampled conformations in each simulation. Prior to clustering, PROTAC conformations were least-squares fitted using all PROTAC atoms to remove overall translational and rotational motion. Pairwise root-mean-square deviations (RMSDs) were calculated using the same atom selection and used for clustering with the “gmx cluster” program in GROMACS 2025.1 [40] with the GROMOS algorithm described by Daura et al. [51]. The RMSD cutoff point of 0.4 nm was selected for the final analysis based on the number and population of clusters and the RMSD distribution. RMSD values were calculated using “gmx rms” and used as a quantitative measure of structural similarity among sampled conformations.

#### 2.4.5 Molecular shape evaluation

The principal moment of inertia (PMI) analysis [52] was performed to describe the global shape of each PROTAC conformer sampled during the simulations. For each trajectory frame, the ligand was selected and its moment of inertia tensor was calculated. Diagonalization of this tensor provided the three principal moments of inertia, (I-1), (I-2), and (I-3), ordered from smallest to largest. To remove the dependence on molecular size, normalized principal moment ratios (NPRs) were calculated as (NPR1 = I-1/I-3) and (NPR2 = I-2/I-3). These descriptors were used to compare the shape distribution of the sampled conformations and to assess whether the molecules preferentially adopted rod-like, disc-like, or more spherical conformational states. Each conformation was classified as rod-like, disc-like/flat, or sphere-like based on its proximity to the idealized normalized PMI vertices at (NPR1, NPR2) = (0, 1), (0.5, 0.5), and (1, 1), respectively.

#### 2.4.6 Simulation analysis

Trajectory analyses, including trajectory preparation, RMSD calculation, feature classification, and RMSD-based clustering, were carried out using in-house Python scripts in combination with GROMACS 2025.1 [40]. Molecular structures, representative conformers, and simulation trajectories were inspected, visualized, and rendered using VMD [53] and PyMOL 3.0.5 [54]. All scripts used for trajectory processing, descriptor calculation, clustering, and downstream analysis are publicly available in the project GitHub repository.

## 3 Results and discussion

### 3.1 PROTAC permeability is independent of conventional molecular descriptors

A set of 20 PROTACs with reported PAMPA permeability values is used to analyze the dynamics of PROTAC molecules, as shown in Figure 1 and summarized in Table 1. The PROTACs were categorized into three permeability classes. PROTAC-1 and PROTAC-2 exhibited markedly high permeabilities (PAMPA *>* 150 nm/s) and were classified as permeable (P). PROTAC-3 showed intermediate-to-high permeability (PAMPA *≈* 14 nm/s). PROTACs with PAMPA values in the range of 2-15 nm/s were classified as moderately permeable (M); and below 2 nm/s were classified as non-permeable (NP).

**Figure 1.**
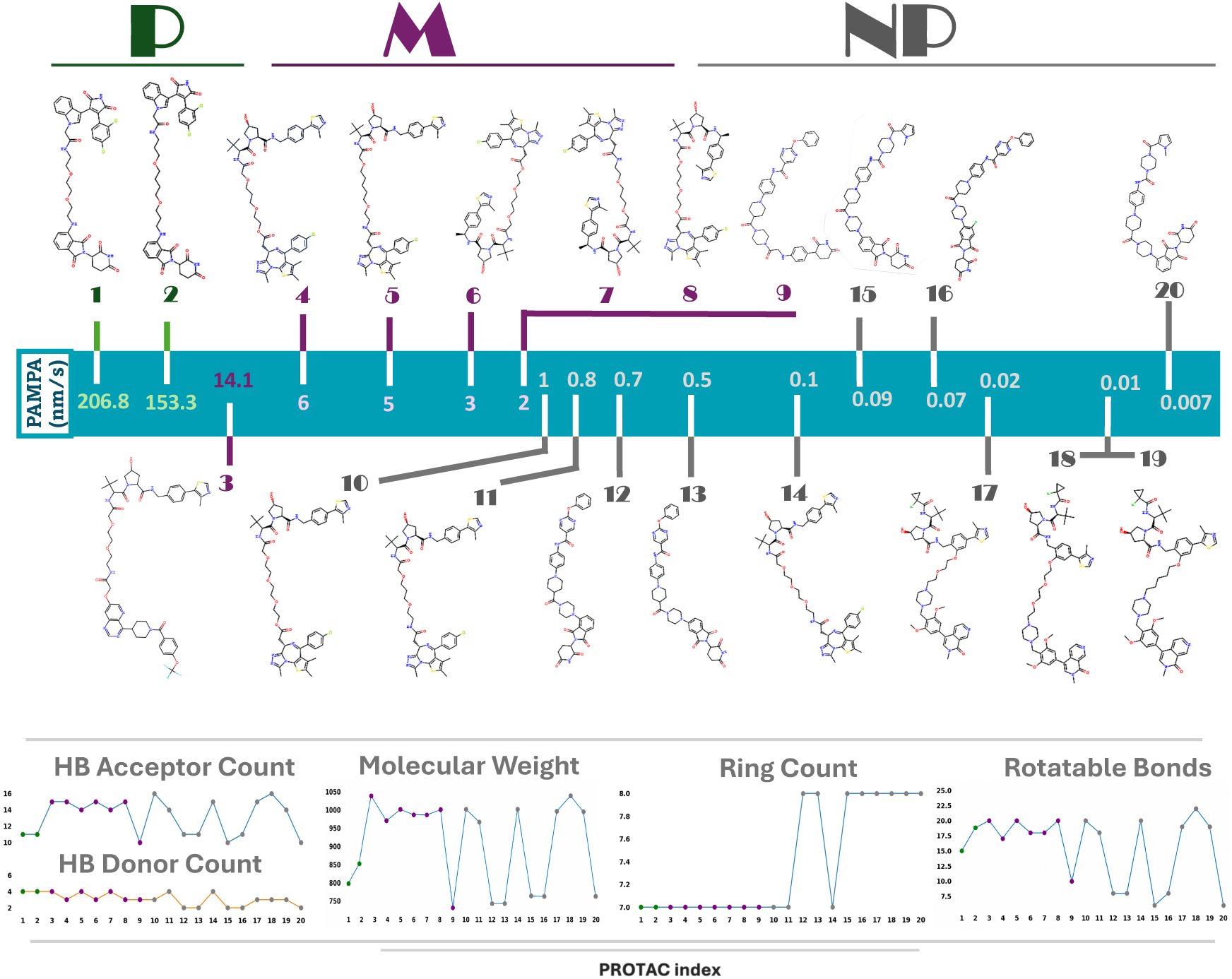
PROTAC permeability is independent of conventional molecular descriptors. The dataset comprises 20 PROTACs spanning a broad range of experimentally measured PAMPA passive permeability values and classified as permeable (P, green), moderately permeable (M, purple), or non-permeable (NP, gray). The corresponding 2D chemical structures are shown relative to their PAMPA permeability values (nm/s). Conventional molecular descriptors, including hydrogen-bond acceptor count, hydrogen-bond donor count, molecular weight, ring count, and rotatable bond count, are shown for each PROTAC in the lower panels. Across the dataset, these descriptors do not exhibit a clear trend with permeability class, indicating that differences in passive permeability cannot be explained by conventional molecular properties alone.

As an initial analysis, conventional 2D molecular descriptors were calculated for each PROTAC to evaluate whether passive permeability could be rationalized by simple properties. These included molecular weight, exact mass, total atom count, heavy atom count, ring count, hydrogen-bond acceptor and donor counts, and rotatable bond count. These descriptors capture only overall molecular size, flexibility, and hydrogen-bonding capacity. They are conformation-independent, and therefore account for neither 3D folding nor intramolecular hydrogen bonding. As shown in Figure 1, PAMPA permeability did not show a clear trend with any of the conventional molecular descriptors examined, indicating PROTAC permeability cannot be explained by simple properties alone and suggesting a stronger role for conformational and solvent-dependent effects.

### 3.2 Enhanced sampling of conformational ensembles reveals that compact permeable-like conformations are not exclusive to highly permeable molecules

Accurate generation of diverse conformational states is particularly important for PROTAC molecules, which are highly flexible and can adopt a broad conformational range, spanning compact folded structures to extended open conformations. However, efficient exploration of this broad conformational space remains challenging, as conventional molecular dynamics simulations may become trapped in local energy minima or require long simulation times to sample relevant conformational transitions. Existing conformer generation approaches, including distance-geometry methods such as RDKit conformer generator [29, 55], knowledge-based sampling [56], force-field-based methods [57], quantum chemical approaches such as Conformer-Rotamer Ensemble Sampling Tool (CREST) [58], and AI-based methods, differ in accuracy, efficiency, and sampling strategies. Although these methods offer specific advantages, balancing conformational diversity, structural correctness, computational cost, and sampling efficiency remains difficult for highly flexible molecules. Moltiverse uses enhanced-sampling molecular dynamics, combining the extended adaptive biasing force (eABF) and metadynamics with the Rg as the collective variable (CV) to promote exploration of both compact and extended conformational states. This strategy enables efficient sampling of diverse conformations across different degrees of molecular compaction while preserving realistic molecular geometries, making it particularly suitable for molecules with many rotatable bonds and complex energy landscapes [34]. For each PROTAC, a conformational ensemble was generated using the Moltiverse protocol, starting from its SMILES representation. The molecule was first converted into a 3D structure and parameterized using the GAFF2 force field. The conformational sampling was carried out in vacuum. Based on atom-count classification, molecules in the medium (72–136 atoms) and medium–large (137–160 atoms) categories were sampled using size-dependent simulation parameters to ensure adequate coverage of conformational space while maintaining computational efficiency. The resulting conformations were subsequently clustered to identify representative structures, yielding 250 clusters per PROTAC. These structures were then refined using molecular mechanics and quantum mechanical optimization to produce a diverse ensemble of low-energy conformers. The generated conformer ensembles were analyzed using in-house Python workflows, through which spatial molecular descriptors were calculated for each conformer. Each conformation was characterized via SASA, and PSA to evaluate molecular polarity, and Rg to quantify size and compactness. IHBs were analyzed in detail, including their number, distances, and angles, to assess their contribution to conformational stability. Additionally, a new descriptor, termed hydrogen-bond ring (HBR) size, was introduced to capture cyclic intramolecular hydrogen-bonding motifs formed when two distant regions of the molecule interact via hydrogen bonding, resulting in a closed ring-like structure; the ring size is defined as the number of atoms enclosed within this cycle. Larger ring sizes were hypothesized to promote intramolecular stabilization and facilitate the formation of compact structures. As shown in Figure 2, the PSA–Rg distribution plot did not reveal a clear separation between permeable and non-permeable PROTACs. Compact conformations with low PSA and low Rg values were observed across all permeability classes, P, M, and NP. This suggests that the ability to adopt compact, less-polar conformations alone is not sufficient to explain the observed differences in passive membrane permeability. Similarly, neither SASA (Supplementary Data and Supplementary Figure S1) nor the intramolecular hydrogen-bonding profiles (Supplementary Data and Supplementary Figure S2) showed a consistent permeability-related trend. Notably, however, PROTAC-1, PROTAC-2, and PROTAC-3 formed a higher number of intramolecular hydrogen bonds than the other compounds (Supplementary Figure S3). Note, however, that the ensembles were generated in vacuum, an environment that favours conformations stabilized by intramolecular H-bonding, so the broad accessibility of compact, low-PSA states across classes is in part expected. Note that they are also cluster representatives rather than weighted trajectory frames, and therefore describe which conformations are geometrically reachable rather than how often each is visited. The comparison in Figure 2 is accordingly a statement about the accessibility of compact states, not about their population. This is an important distinction to make for the solvated simulations below. This suggests that permeability may depend more on the stability and population of certain conformations, and solvent-dependent behavior rather than ability to form compact structures. Therefore, in the next step, selected compact conformers were evaluated *in silico* in different solution environments.

**Figure 2.**
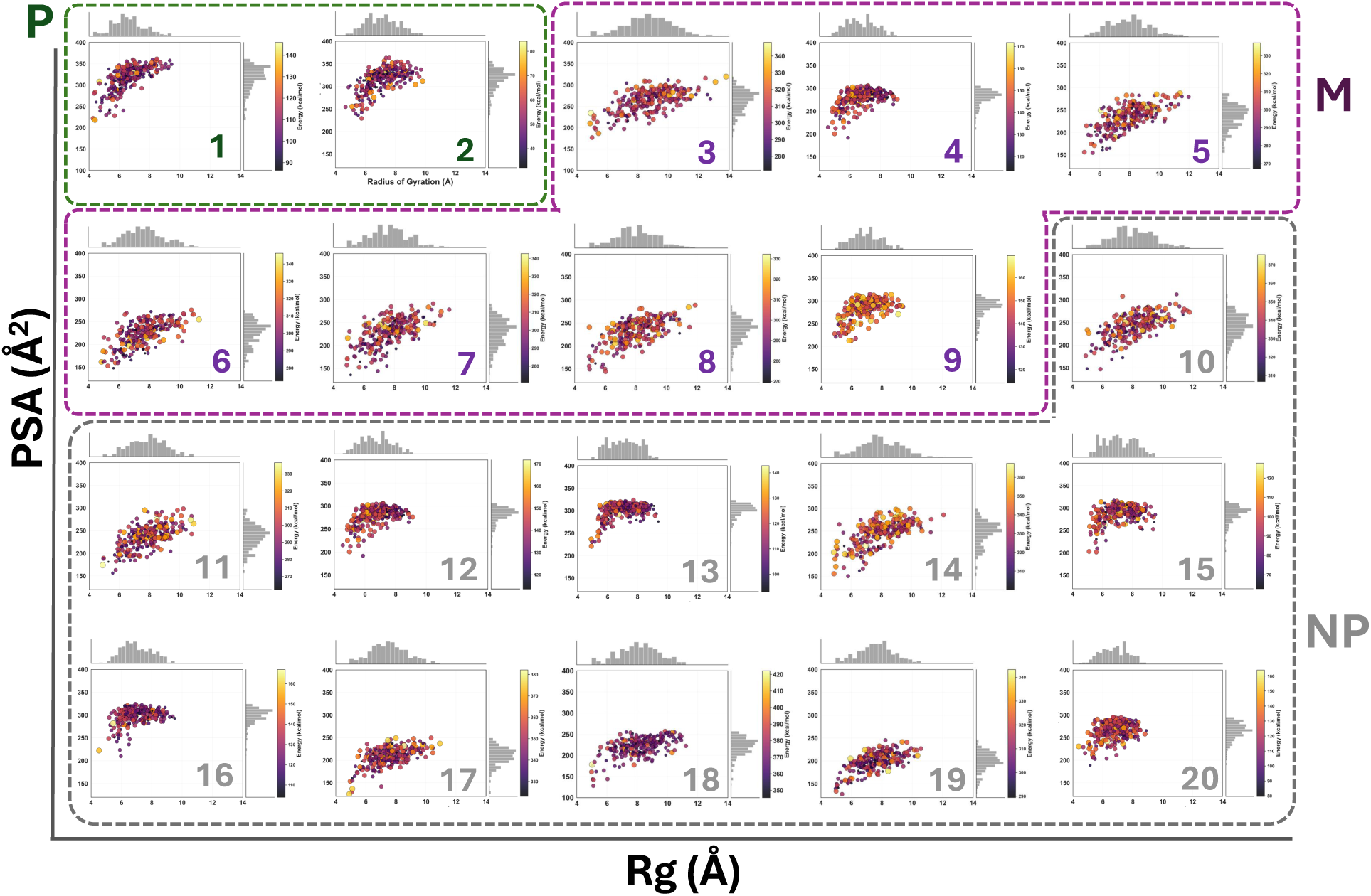
Enhanced sampling of conformational ensembles reveals that compact permeable-like conformations are not exclusive to highly permeable molecules. 2D distributions of radius of gyration (Rg) and polar surface area (PSA) for conformational ensembles generated by Moltiverse enhanced-sampling molecular dynamics (ES-MD) for the 20-PROTAC dataset. Each point corresponds to a sampled conformer, with color and size reflecting its energy. Energy values indicate the relative energetic distribution of conformers within each PROTAC ensemble and should not be directly compared between different PROTAC molecules. The corresponding marginal distributions of Rg and PSA are shown along the axes. PROTACs are grouped according to their experimental PAMPA permeability classification as permeable (P, green), moderately permeable (M, purple), and non-permeable (NP, gray). All three permeability classes access compact conformations characterized by low Rg and PSA, demonstrating that the ability to adopt compact, low-polar-surface-area states is not exclusive to highly permeable PROTACs and, by itself, does not explain differences in passive permeability.

### 3.3 Maximum hydrogen bond ring size (MRS) is used to select comparable compact states across passive permeability ranges of PROTACs

Representative compact conformations were selected from three classes: highly permeable PROTAC-1 (P, 206.8 nm/s) [30], moderately permeable PROTAC-3 (M, 14.1 nm/s)[10], and non-permeable PROTAC-18 (NP, 0.01 nm/s)[33], based on the maximum ring size (MRS) metrics. MRS was defined as the size of the largest HBR and used as an indicator of long-range intramolecular hydrogen bonding and molecular folding. A related geometric descriptor, SMID, which represents the smallest effective molecular diameter, was introduced by Bower et al. [59] as part of the BBI index [18]. Because SMID and Rg are both purely geometric, neither identifies the intramolecular interactions that stabilize a folded conformation. Although MRS showed no relationship with PAMPA permeability across the 20 ensembles, consistent with the other descriptors examined, it provided a criterion for identifying folded conformers whose compactness is supported by a long-range intramolecular hydrogen bond rather than inferred from geometry alone. The selected compact conformers of PROTAC-1, PROTAC-3, and PROTAC-18 contained 10, 5, and 6 IHBs, respectively, with MRS of 27, 25, and 34 atoms. Notably, PROTAC-1 contained four IHBs forming large pseudo-rings, whereas PROTAC-3 and PROTAC-18 each contained only one such long-range interaction. Despite different permeability profiles, the selected conformers adopted similar compact structures, as indicated by their comparable Rg, PSA, and SASA values. PROTAC-1 had an Rg of 5.69 Å, SASA of 996.94 Å², and PSA of 228.10 Å², while PROTAC-3 showed an Rg of 5.17 Å, SASA of 984.14 Å², and PSA of 199.84 Å². The non-permeable PROTAC-18 also displayed a compact structure, with an Rg of 5.05 Å, SASA of 1022.06 Å², and PSA of 143.81 Å² (Figure 3). These selected compact conformers were used as the initial structures for subsequent MD simulations. The initial conformations of the three representative systems showed distinct positions in the normalized principal moment of inertia (PMI) space. The P initial structure had NPR1 and NPR2 values of 0.42 and 0.67, respectively, and was classified as disc-like shape, indicating a relatively extended and flattened molecular geometry. In contrast, both the M and NP initial structures were classified as sphere-like, with NPR1/NPR2 values of 0.79/0.90 and 0.70/0.82, respectively. The geometric descriptors indicate more compact and isotropic starting geometries compared with the P structure.

**Figure 3.**
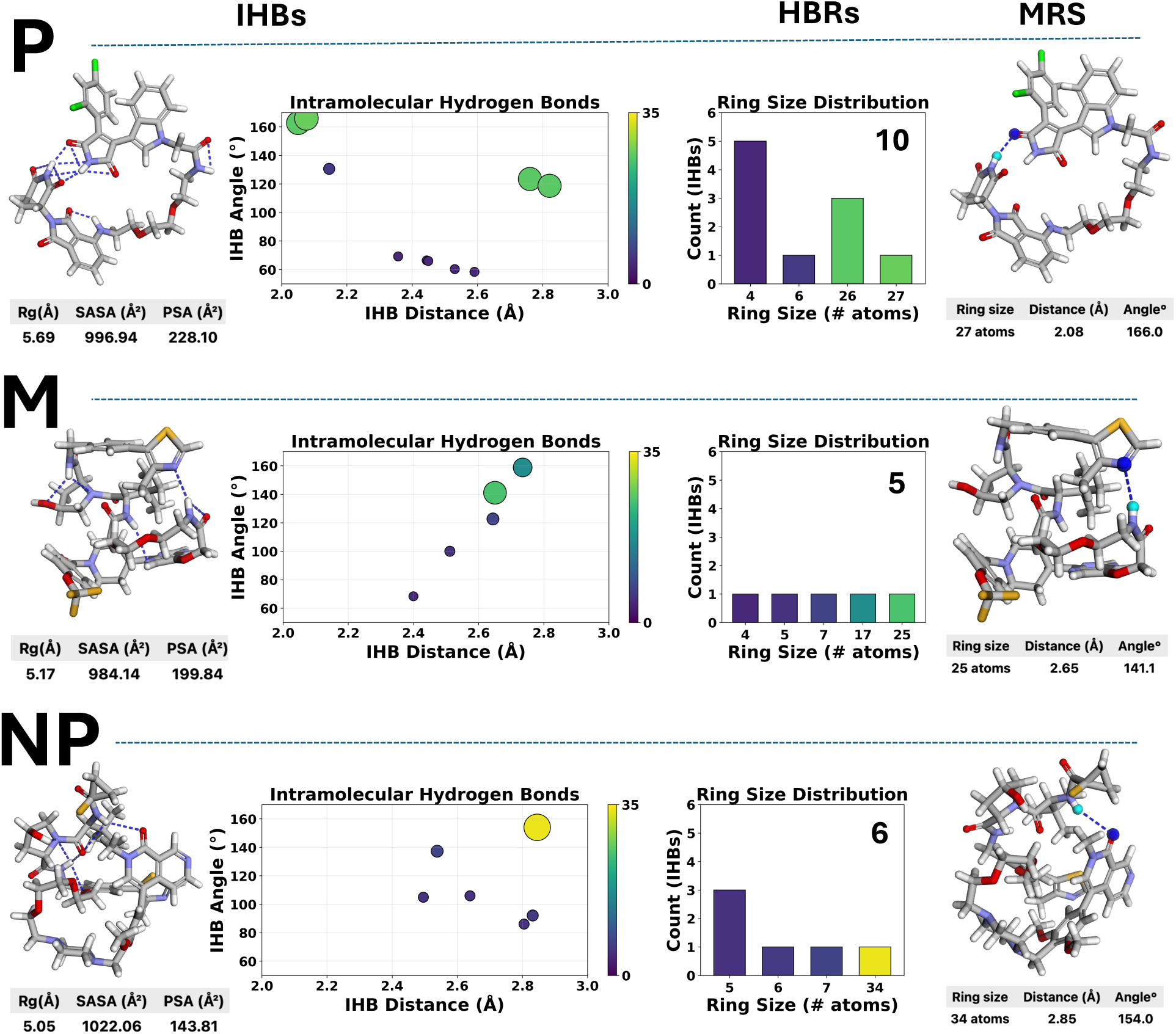
Maximum hydrogen-bond ring size (MRS) is used as a structural criterion to identify compact conformations stabilized by long-range interactions and select representative compact states across PROTAC permeability classes. Representative compact conformations of highly permeable PROTAC-1 (P), moderately permeable PROTAC-3 (M), and non-permeable PROTAC-18 (NP) were selected and characterized based on their intramolecular hydrogen-bonding (IHB) patterns. Left: compact structures with all identified IHBs shown as blue dashed lines, together with their radius of gyration (Rg), solvent-accessible surface area (SASA), and polar surface area (PSA). Middle: IHB patterns are characterized by the total number of bonds, their H···acceptor distance and hydrogen-bond angles, and the ring size associated with each interaction. Right: the IHB forming the MRS is highlighted for each representative structure, with its ring size (number of atoms), H···acceptor distance, and hydrogen-bond angle reported.

### 3.4 Stability of permeable-like compact states evaluated via U-MD simulations in polar and apolar environments

To evaluate the stability of the selected folded conformers, all-atom U-MD simulations were performed in two solvent environments; water and chloroform. Chloroform is commonly used as a low-polarity membrane-mimetic solvent and the approximate dielectric constant of the lipid-bilayer interior [60].

In contrast, water provides a highly polar medium that is more representative of physiological conditions, allowing hydrophilic interactions and solvent-exposed molecular features to be evaluated. In total, six simulation systems were prepared, corresponding to three representative PROTACs 1, 3, and 18, simulated in each of the two solvents. For each system, four independent replicas of 200 ns were performed, resulting in a total simulation time of 800 ns per system. The structural dynamics of conformers were first evaluated by calculating the RMSD relative to their initial structure, to quantify the overall deviation during the simulations. PROTAC-1 showed the lowest RMSD in the apolar environment, 2.18 ± 0.51 Å, indicating the strongest preservation of the starting structure. Its RMSD increased to 4.53 ± 0.25 Å in the polar environment, suggesting greater structural rearrangement in water despite the relatively compact Rg distribution. PROTAC-3 showed similar RMSD values in both solvents, 6.86 ± 0.67 Å in the apolar environment and 6.87 ± 1.29 Å in the polar environment. PROTAC-18 showed an RMSD of 6.03 ± 0.93 Å in the apolar environment and 4.75 ± 1.02 Å in the polar environment, indicating better preservation of the compact state in water than in chloroform. Overall, the RMSD analysis indicates compound-dependent solvent effects, with PROTAC-1 folded conformer being stable in chloroform and water, and PROTAC-18 showing greater stability in water (Figure 4 and supplementary Figure S4)

**Figure 4.**
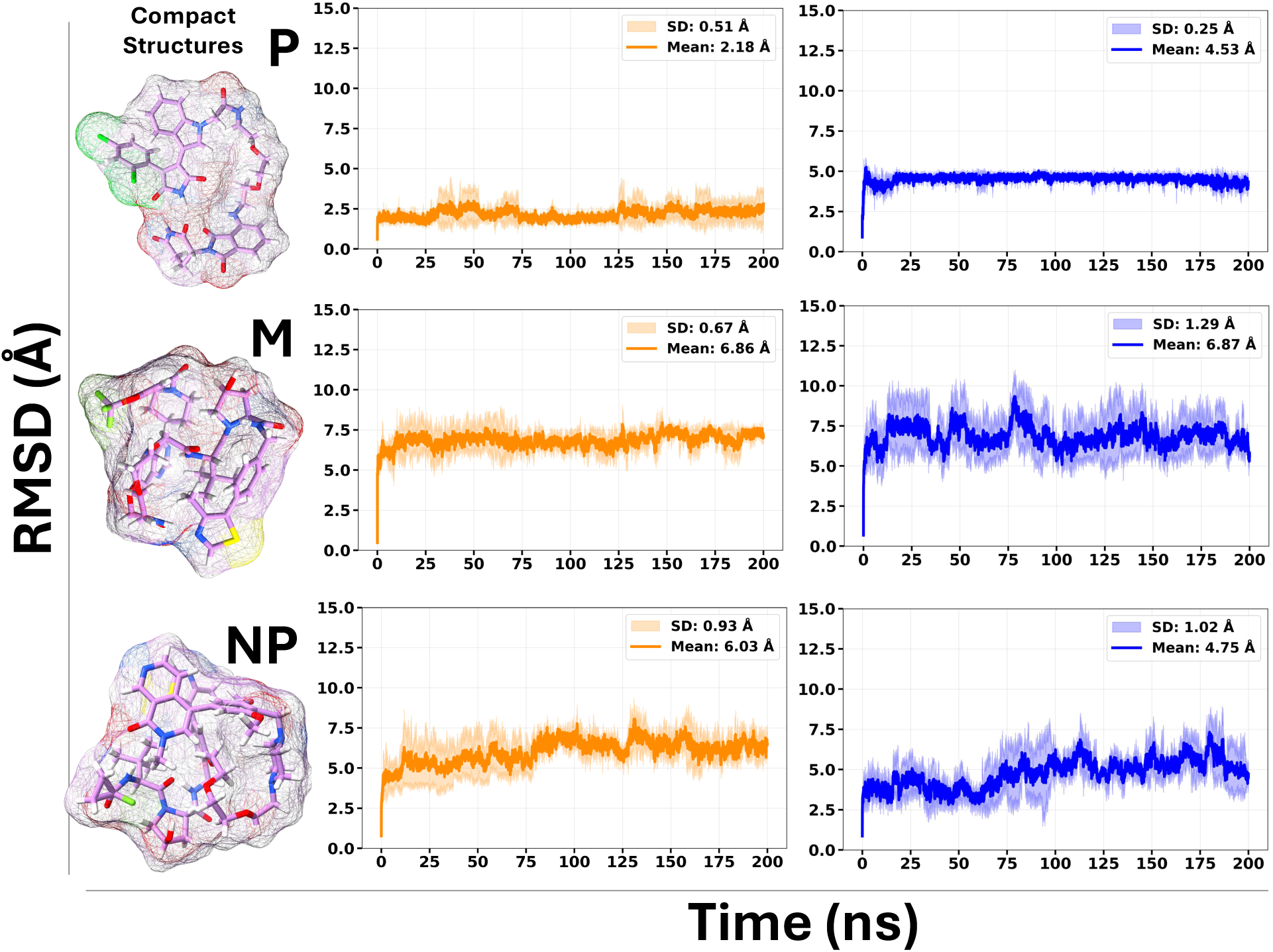
Solvent-dependent conformational stability of PROTACs initiated from compact structures. Root-mean-square deviation (RMSD) profiles of highly permeable PROTAC-1 (P), moderately permeable PROTAC-3 (M), and non-permeable PROTAC-18 (NP) during all-atom MD simulations in chloroform (orange) and water (blue). For each system, four independent 200ns replicas were performed, corresponding to a total sampling time of 800 ns per system. RMSD was calculated relative to the corresponding compact initial structure. Solid lines represent the mean RMSD across the four replicas, and shaded margins indicate the time-averaged standard deviation (SD). The compact conformation is shown on the left as a stick representation overlaid with a mesh molecular surface.

As shown in Figure 5, The Rg and PSA distributions were analyzed to characterize changes in molecular compactness and polar surface exposure throughout the simulations. In the ES-MD distributions, the red markers denote the compact conformers selected as the initial structures for the subsequent U-MD simulations. For PROTAC-1, the initial compact structure had an Rg of 5.69 Å and a PSA of 228.10 Å². In the apolar environment, the Rg mean/mode was 5.65/5.64 Å, with a range of 4.55–8.22 Å, indicating that PROTAC-1 remained close to its initial compact state. In the polar environment, the Rg mean/mode decreased slightly to 5.00/4.97 Å, with a similar range of 4.52–8.15 Å, suggesting that the molecule maintained a compact conformation in water. The PSA mean/mode was 194.98/192.38 Å² in the apolar environment and 225.02/236.04 Å² in the polar environment. Compared with the initial PSA, PROTAC-1 showed lower polar surface exposure in chloroform, while the polar simulation remained closer to the initial PSA value.(Figure 5, and supplementary Figure S5.b).

**Figure 5.**
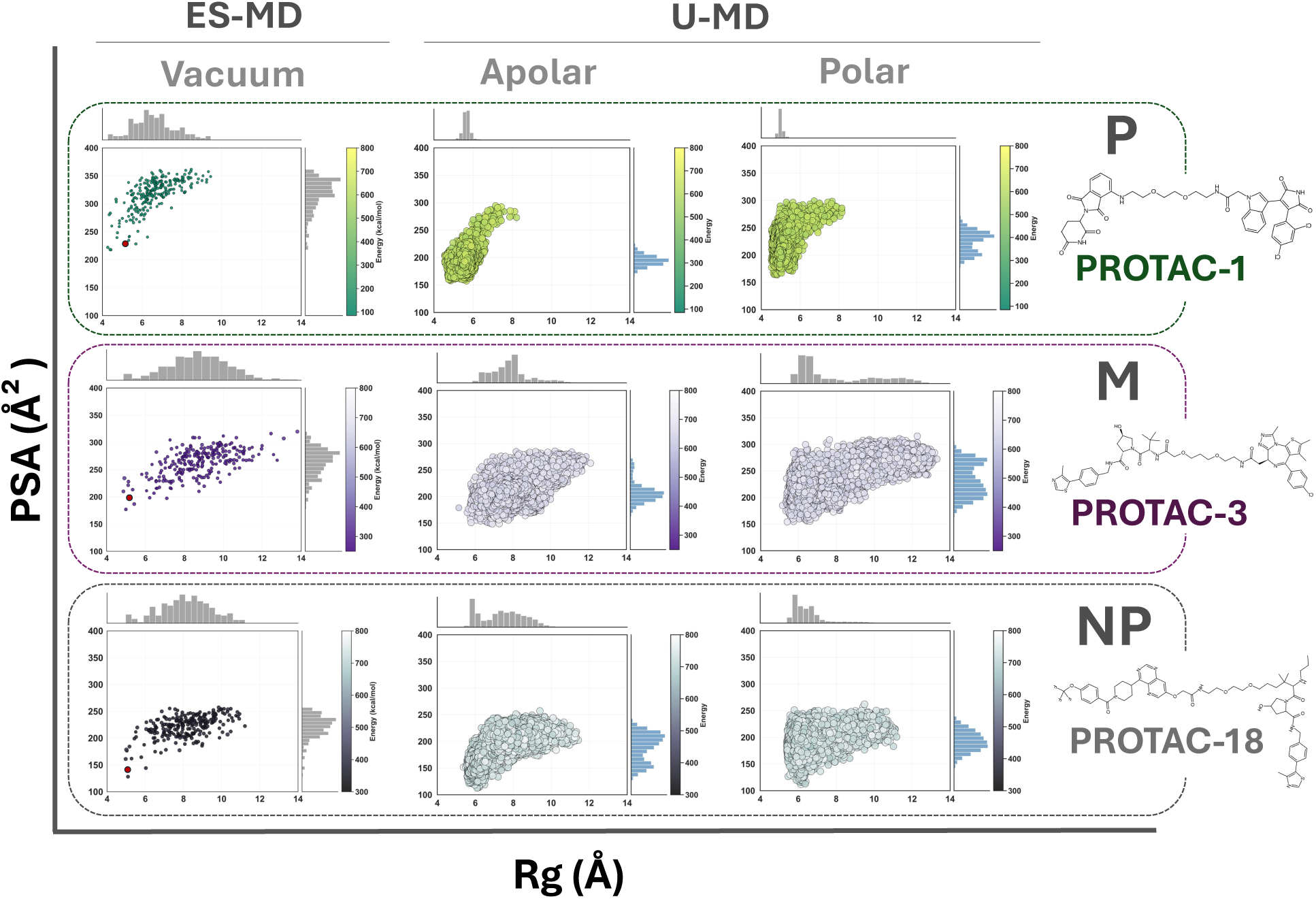
Stability of compact states evaluated via unbiased MD simulations in polar and apolar Environments. Conformational ensembles of highly permeable PROTAC-1 (P), moderately permeable PROTAC-3 (M), and non-permeable PROTAC-18 (NP) are represented by their radius of gyration (Rg) and polar surface area (PSA). Left: ES-MD ensembles, where the red dot indicates the compact conformation selected using MRS to initiate subsequent all-atom U-MD simulations. Middle and right: conformational ensembles sampled during U-MD in apolar (chloroform) and polar (water) environments, respectively, showing the structural changes from the selected compact starting state. Each point represents a sampled conformation, with both point color and size indicating its energy. Energy values are interpreted only within the ensemble of each individual PROTAC and are not directly compared between different molecules. Marginal histograms show the distributions of Rg and PSA.

For PROTAC-3, the initial compact structure had an Rg of 5.17 Å and a PSA of 199.84 Å². In contrast to PROTAC-1, PROTAC-3 showed a clear increase in Rg during the simulations. In the apolar environment, the Rg mean/mode increased to 7.87/8.10 Å, with a range of 5.14–12.10 Å. In the polar environment, the Rg mean/mode was 7.97/6.50 Å, with a wider range of 5.16–13.32 Å. These values indicate that PROTAC-3 moved away from the initial compact structure in both solvents. The PSA increased to mean/mode of 207.79/201.30 Å² in the apolar environment and 233.75/271.10 Å² in the polar environment, indicating slightly greater polar surface exposure in water. For PROTAC-18, the initial compact structure had an Rg of 5.05 Å and a PSA of 143.81 Å². In the apolar environment, the Rg mean/mode increased to 7.41/5.77 Å, with a range of 5.39–11.42 Å. In the polar environment, the Rg mean/mode was lower, 6.57/5.77 Å, with a range of 5.40–11.18 Å. These values show that PROTAC-18 deviated from its initial compact state in both solvents, but the increase in Rg was smaller in the polar environment. The PSA mean/mode increased relative to the initial value in both solvents, reaching 188.72/208.72 Å² in the apolar environment and 192.74/183.37 Å² in the polar environment. Overall, PROTAC-1 remained the most stable relative to its initial compact structure, especially in the apolar environment. PROTAC-3 showed the largest structural rearrangement in both environments, as reflected by high RMSD values and increased Rg. PROTAC-18 showed intermediate behavior, with greater deviation in chloroform but comparatively preserved compact-state in water. Across the three compounds, the polar environment generally resulted in lower or more compact Rg behavior for PROTAC-1 and PROTAC-18, while PSA tended to increase in water for PROTAC-1 and PROTAC-3 (Table 2 and Supplementary Figure S5.b)

**Table 2.** Structural properties of PROTAC-1, PROTAC-3, and PROTAC-18 during U-MD simulations in polar (water) and apolar (chloroform) environments, calculated over the concatenated 800 ns ensemble. For RMSD, the reported SD is the time-averaged SD across four replicas at corresponding time points.

| Class | Solvent environment | RMSD vs initial structure (SD)(Å) | Initial Rg (Å) | Rg mean/mode (min–max) (Å) | Initial PSA (Å<sup>2</sup>) | PSA mean/mode (min–max) (Å<sup>2</sup>) |
| --- | --- | --- | --- | --- | --- | --- |
| **P** | Apolar | 2.18 (0.51) | 5.69 | 5.65/5.64<br>(4.55–8.22) | 228.10 | 194.98/192.38<br>(157.47–294.53) |
|  | Polar | 4.53 (0.25) |  | 5.00/4.97<br>(4.52–8.15) |  | 225.02/236.04<br>(163.21–299.33) |
| **M** | Apolar | 6.86 (0.67) | 5.17 | 7.87/8.10<br>(5.14–12.10) | 199.84 | 207.79/201.30<br>(148.47–285.70) |
|  | Polar | 6.87 (1.29) |  | 7.97/6.50<br>(5.16–13.32) |  | 233.75/271.10<br>(154.35–315.38) |
| **NP** | Apolar | 6.03 (0.93) | 5.05 | 7.41/5.77<br>(5.39–11.42) | 143.81 | 188.72/208.72<br>(112.53–257.18) |
|  | Polar | 4.75 (1.02) |  | 6.57/5.77<br>(5.40–11.18) |  | 192.74/183.37<br>(112.16–262.98) |

Although the U-MD simulations were initiated from selected compact conformers and were not designed for exhaustive conformational sampling, their Rg–PSA (Supplementary Figure S5.a) and Rg-SASA distributions (Supplementary Figure S1) provided a useful comparison with the MV-generated ensembles. As expected, the U-MD distributions differed from the broader Moltiverse distributions and did not show the same balanced coverage of conformational space. Nevertheless, the simulated structures partially overlapped with regions sampled by Moltiverse, particularly along the Rg dimension. This overlap indicates that the selected Moltiverse conformers represent physically accessible compact and intermediate states that can be reached or maintained in solvent.

### 3.5 Solvent-dependent structural dynamics reveal dominant PROTAC conformational states

Structural analyses were then performed to evaluate whether the initially closed structures remained stable or rearranged into alternative, more extended conformations in a solvent-dependent manner. First, PMI analysis was performed to characterize the global shape of the sampled ensemble. The P system starts from a more flattened conformation, whereas the M and NP systems begin from sphere-like conformations. In the apolar environment, PROTAC-1 predominantly retained a disc-like/flat shape similar to its initial conformation, with 86.8% of the sampled structures classified in the disc-like/flat region and only 13.0% classified as rod-like. The minor rod-like population likely reflects transient opening event deviations during the simulation, after which the molecule returned to the dominant and more stable disc-like/flat state. A subsampled trajectory and the corresponding movie are also available in the project’s Zenodo repository. In contrast, PROTAC-3 and PROTAC-18 moved away from their starting shapes in both solvent environments. PROTAC-3 shifted predominantly toward rod-like conformations in both environments, with rod-like structures accounting for 83.0% and 66.4% of the sampled conformers in the apolar and polar simulations, respectively. PROTAC-18 also shifted toward rod-like conformations in the apolar environment, where 68.0% of the structures were classified as rod-like and 31.4% as disc-like/flat. In the polar environment, however, PROTAC-18 sampled a predominantly disc-like/flat ensemble, rising to 61.9% of conformers classified as disc-like/flat and 35.8% as rod-like (Figure 6 and Supplementary Figure S6). The comparison between different classes demonstrates that shape alone cannot fully explain permeability.

**Figure 6.**
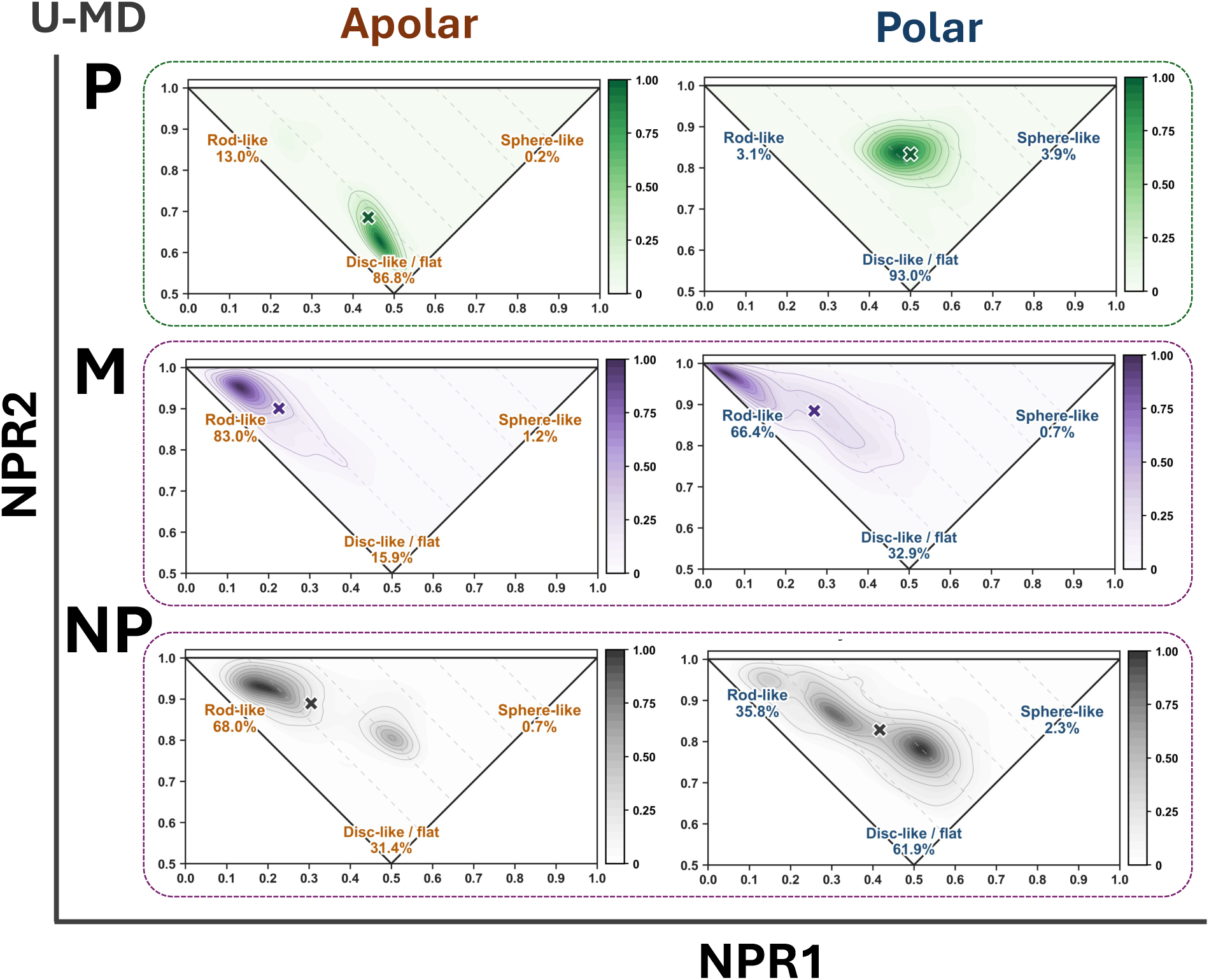
PMI-based shape distributions of permeable (P), moderately permeable (M), and non-permeable (NP) PROTACs during U-MD simulations in apolar and polar environments. Percentages indicate the rod-like, disc-like/flat, and sphere-like populations; red circles mark the initial compact structures, and crosses indicate the ensemble means.

Then, RMSD-based clustering was performed to identify the dominant conformational states sampled by each PROTAC using gmx cluster and RMSD cutoff of 4 Å. Only the three most populated clusters are reported, while other clusters contributing less than 5% of the trajectory were excluded from the analysis and treated as low-population noise. The results revealed clear differences in conformational heterogeneity among the three compounds. In the apolar environment, PROTAC-1 was almost exclusively represented by a compact conformational state, accounting for 99.20% of the trajectory. In water, the molecule adopted a different closed arrangement, which was similarly dominant and represented 98.50% of the sampled structures. In contrast to PROTAC-1, PROTAC-3 displayed a more heterogeneous conformational ensemble in both solvent environments. In the apolar simulation, the three most populated clusters accounted for 42.31%, 16.29%, and 14.49% of the trajectory, while in water the corresponding clusters represented 33.63%, 18.23%, and 9.61% of the sampled structures. The absence of a highly populated single cluster indicates that PROTAC-3 sampled multiple conformational states rather than remaining confined to one dominant arrangement. For PROTAC-18, the clustering analysis also indicated solvent-dependent conformational behavior. In the apolar environment, the trajectory was distributed across several populated clusters, with the three largest clusters accounting for 29.57%, 23.15%, and 16.89% of the sampled structures. In contrast, the polar simulation showed a more dominant cluster, accounting for 62.64% of the trajectory, followed by smaller clusters of 17.19% and 6.09%. As expected, given that both measure deviation from the starting structure, the clustering results track the trajectory RMSD profiles over time (Figure 4), which showed smaller structural deviations for PROTAC-1 in the apolar (2.18 ± 0.51 Å) and polar (4.53 ± 0.25 Å) and for PROTAC-18 in the polar environment (4.75 ± 1.02 Å). Figure 7 presents the PMI-based shape classification and the corresponding PSA and Rg values for the “representative structures” of the most populated RMSD-based clusters in each ensemble.

**Figure 7.**
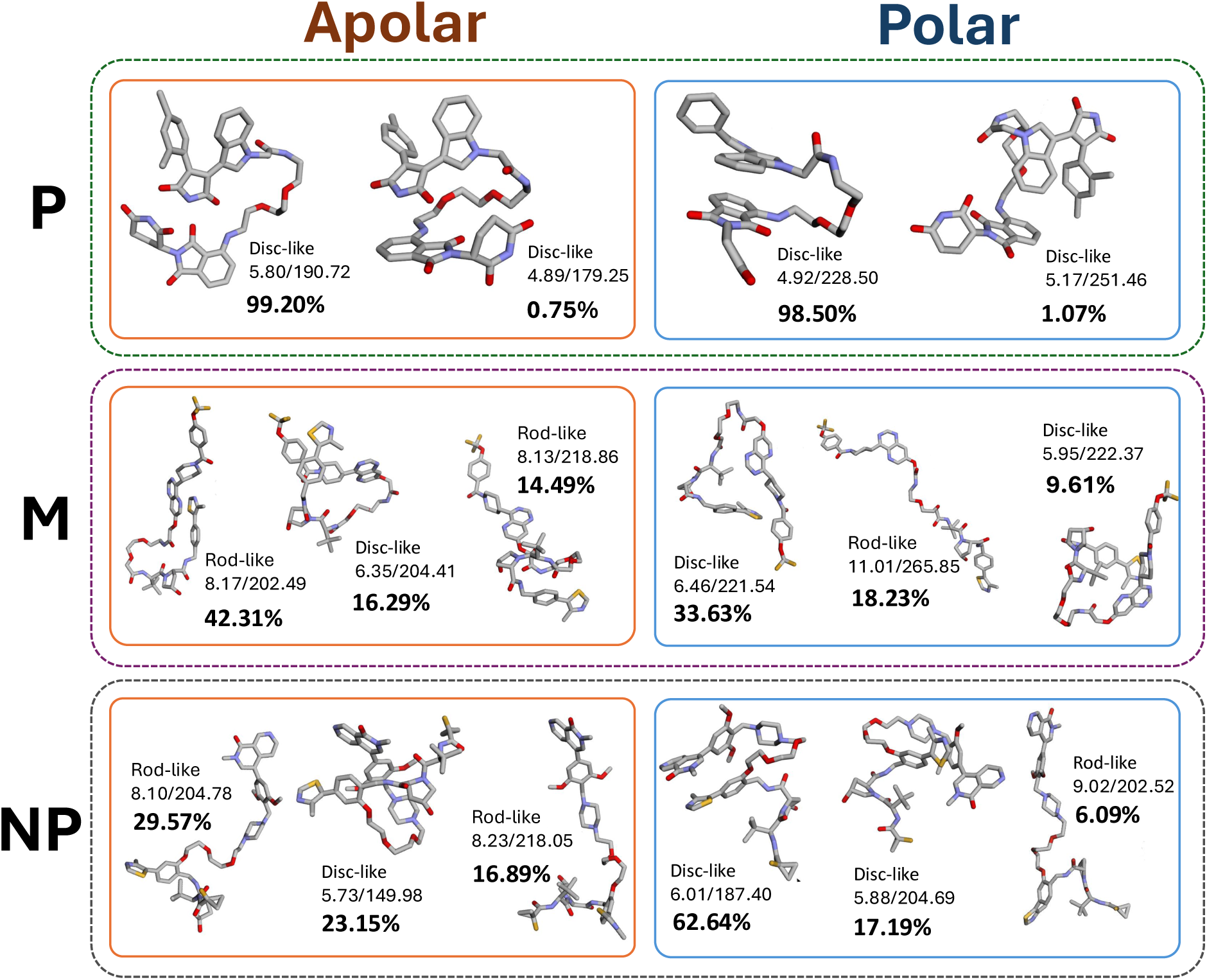
Representative structures of the dominant clusters identified by RMSD-based clustering during U-MD simulations of permeable (P), moderately permeable (M), and non-permeable (NP) PROTACs in apolar and polar environments. Labels indicate shape, Rg/PSA (Å/Å²), and cluster population.

For comparison purposes, the clusters were merged according to their molecular shape using an Rg threshold of 6.5 Å to distinguish closed and open conformations, and PMI analysis to classify the structures as flat/disc-like or rod-like. Weighted Rg and PSA values were calculated to provide structural descriptors of each conformational ensemble (Table 3). The descriptor values calculated for the representative structure of the dominant cluster showed minor differences from the corresponding population-weighted values obtained from all members of the simulation ensemble; however, the overall trends remained consistent. The permeable PROTAC-1 showed the most stable conformational behavior in both solvent environments. In the apolar environment, the dominant ensemble remained highly compact and closely related to the initial state, with a weighted Rg of 5.80 Å and a weighted PSA of 190.66 Å². Almost the entire clustered population (99.99%) was classified as closed-flat, with no appreciable open-rod population detected. This indicates that the molecule did not shift away from its starting shape and size. A similar compact behavior was observed in the polar environment. Although the weighted PSA increased to 228.83 Å², the weighted Rg remained low (4.93 Å), and 99.58% of the ensemble was still classified as closed-flat. This suggests that PROTAC-1 preserved a compact flat conformational state in both solvents. However, the polar closed-flat state likely represents a different compact arrangement from that observed in the apolar environment, suggesting that the molecule adjusts its polar surface exposure according to the surrounding solvent. Notably, MRS-selected conformers of PROTAC-1 remained stable during the subsequent solvent simulations, although with one compound per class this indicates only that the criterion can identify persistent folds, not that high MRS implies persistence. In contrast, the moderately permeable PROTAC-3 displayed substantially greater conformational flexibility. In the apolar environment, the weighted Rg increased to 7.92 Å, indicating a more extended ensemble than the permeable compound. The dominant conformational population shifted towards open-rod structures, accounting for 69.35% of the clustered conformations, whereas only 16.30% remained closed-flat. In the polar environment, the weighted Rg remained similarly high (7.79 Å) and the conformational ensemble became more heterogeneous, containing 43.25% closed-flat and 32.23% open-rod conformations. The weighted PSA showed only a modest decrease in the apolar environment (230.54 → 209.82 Å²), suggesting partial shielding of polar groups. The non-permeable PROTAC-18 exhibited a different solvent-dependent behavior. In the polar environment, it remained relatively compact, with a weighted Rg of 6.22 Å, a relatively low weighted PSA of 193.29 Å², and a dominant closed-flat population (83.08%). However, upon transfer to the apolar environment, the conformational ensemble became more heterogeneous. The weighted Rg increased to 7.23 Å, while the dominant population shifted towards open-rod conformations (53.33%), although a substantial fraction (30.31%) remained closed-flat. Interestingly, the weighted PSA slightly decreased in the apolar environment (193.29 → 187.18 Å²), indicating some reduction in exposed polarity despite the increased conformational extension. Taken together, the permeable PROTAC remained almost exclusively closed-flat in both solvents and showed a reduction in PSA under apolar conditions. In contrast, both the moderately permeable and non-permeable compounds populate substantially larger fractions of open-rod conformations in the apolar environment, indicating weaker preservation of the initial compact state. Despite being classified as non-permeable, PROTAC-18 retains a larger closed-flat population in the apolar environment (30.31%) than PROTAC-3 (16.30%) and also exhibits a lower weighted PSA (187.18 Å² versus 209.82 Å²). Our results suggest that molecular compactness and shape are not sufficient to distinguish permeable from non-permeable, at least within the limited timescale and conformational sampling of the present U-MD simulations.

**Table 3.** Population-weighted Rg and PSA values and the distributions of closed-flat and open-rod conformations for each U-MD conformational ensemble in apolar and polar environments. Only closed-flat and open-rod are reported; the remainder comprises closed-rod, open-flat, and sphere-like conformations. Populations exceeding 80% are shown in bold. Full details are provided in the supplementary Excel file.

| Class | Solvent | Weighted Rg (Å) | Weighted PSA (Å<sup>2</sup>) | Closed-flat (%) | Open-rod (%) | Others (%) |
| --- | --- | --- | --- | --- | --- | --- |
| **P** | Apolar | 5.80 | 190.66 | **99.99** | 0.00 | 0.01 |
|  | Polar | 4.93 | 228.83 | **99.58** | 0.42 | 0.00 |
| **M** | Apolar | 7.92 | 209.82 | 16.30 | 69.35 | 14.35 |
|  | Polar | 7.79 | 230.54 | 43.25 | 32.23 | 24.52 |
| **NP** | Apolar | 7.23 | 187.18 | 30.31 | 53.33 | 16.36 |
|  | Polar | 6.22 | 193.29 | **83.08** | 9.32 | 7.60 |

What does separate the three compounds is not whether a compact state is accessible, but whether one is retained. PROTAC-1 remained in a closed-flat ensemble in both solvents (99.99% and 99.58%) of the clustered population, yet the apolar and polar ensembles correspond to structurally distinct arrangements. The molecule therefore does not simply remain folded; it exchanges between two well-defined folded states as the environment changes, which is the behavior the chameleonicity hypothesis predicts. Among these three compounds, PROTAC-1 is the only one that persists across a change of environment. By contrast, PROTAC-3 did not retain its fold in either environment, and PROTAC-18 lost its fold in chloroform while retaining a predominantly closed-flat ensemble in water (83.08%).

Compared with the NMR-derived ensembles reported in study [10], our simulations showed a different solvent response for PROTAC-3. The NMR study identified a more compact and less polar ensemble in apolar than in polar environment, with population-weighted Rg/PSA values of 5.93 Å/200 Å² and 6.93 Å/226 Å², respectively, supporting solvent-dependent conformational adaptation. Our results indicate little solvent-dependent change in overall size, but the PSA values were different in apolar and polar conditions. The dominant conformations also differed: in chloroform, our most populated cluster was an open-rod-like state with a population of 42.31%, Rg = 8.17 Å, and PSA = 202.49 Å², whereas in water the dominant cluster was a more compact, closed-flat state with a population of 33.63%, Rg = 6.46 Å, and PSA = 221.54 Å². Thus, PROTAC-3 exhibited solvent-dependent redistribution among conformational states, but not the same global compaction pattern observed in the NMR-derived ensembles. Similar discrepancies between simulation- and NMR-derived ensembles have been previously reported by Poongavanam et al. [12], with densely populated regions identified by MD exhibiting larger Rg values than the corresponding conformational populations reconstructed using NAMFIS algorithm (NMR analysis of molecular flexibility in solution) [61]. In our study, this discrepancy may also partly reflect the limited MD timescale in our study and possible bias introduced by initiating the simulations from a compact structure stabilized by strong intramolecular hydrogen bonds, whereas the experimental study obtained broader solution ensembles constrained by NMR data. In addition, the solvent conditions were not identical: our simulations used chloroform (CHCl3) and pure water, whereas the NMR experiments used deuterated chloroform (CDCl3) and DMSO-d6–D2O (10:1). The latter difference is particularly important because DMSO and water can produce markedly different conformational ensembles as reported by Bhusal and Valsson [62]. They examined MZ1 (PROTAC-14) in chloroform, water, and DMSO using enhanced-sampling atomistic simulations based on parallel-bias metadynamics, in which all rotatable dihedral angles were biased as local collective variables and found that MZ1 preferentially adopted compact, folded conformations in both water and chloroform, whereas substantially more extended conformations were populated in DMSO. Therefore, the experimental NMR DMSO-d6–D2O mixture may not be directly comparable to the water environment used in our simulations, and could partly contribute to the observed disagreement.

Previously, Abeje et al. [11] investigated nine closely related VHL-based PROTACs and demonstrated that relatively small changes in linker composition can markedly affect membrane permeability. Based on NMR measurements in chloroform and conformational ensembles reconstructed using the NAMFIS algorithm, they proposed that higher permeability is associated with the ability to adopt folded conformations with lower solvent-accessible polar surface area and molecular dimensions in low-polarity environments. Nevertheless, despite the pronounced difference in permeability, the ensemble-weighted structural descriptors differed only moderately. The highly permeable PROTAC (38.02 nm/s) exhibited a PSA of 170.9 Å², and an Rg of 6.46 Å, whereas the medium–low permeable PROTAC (0.52 nm/s) showed corresponding values of 206.3 Å² and 6.98 Å. Thus, although the authors conclude that linker-dependent folding helps explain the observed permeability, the NMR comparison is based on only two ensembles and an Rg difference of 0.52 Å (within the variance observed for that property), limiting the quantitative discriminatory power of these descriptors. A similar trend was reported by Poongavanam et al. [12] for three closely related CRBN-based PROTACs. In contrast, our study showed that PROTACs from all permeability classes could adopt compact conformations during both ES-MD and apolar U-MD simulations, although these states differed in their populations and persistence.

## 4 Limitations

The dataset is small and unevenly distributed across permeability classes. The permeable class contains only two compounds, PROTAC-1 and PROTAC-2, which come from the same study [30] and share an identical TPSA, ring count, HBA and HBD, differing by four heavy atoms and four rotatable bonds. Conclusions about permeable PROTACs therefore rest on a single chemical series. The PAMPA values were also compiled from different literature sources, which differ in experimental protocol, reporting, and measurement uncertainty. The two permeable compounds were measured by BBB-PAMPA and the remaining 18 by GIT-PAMPA, so the permeable class also differs from the others in assay type. Values from different PAMPA models and laboratories are not directly comparable, and the class boundaries should therefore be read as an ordinal grouping rather than as calibrated thresholds. We generated the ES-MD ensembles in vacuum, and we simulated only one representative compound per class in solution. The three differ in size and flexibility as well as in permeability: PROTAC-1 is the smallest and least flexible (801 Da, 56 heavy atoms, 15 rotatable bonds), while PROTAC-3 and PROTAC-18 are considerably larger (1034 and 1041 Da, 73 and 74 heavy atoms, 20 and 22 rotatable bonds). The greater conformational persistence of PROTAC-1 may reflect its smaller size rather than its permeability class, and separating the two effects would require a size-matched permeable and non-permeable pair. PROTAC-3 and PROTAC-18 are themselves closely matched in size and flexibility yet differ by more than three orders of magnitude in PAMPA permeability and behave differently in solution, so size does not account for the differences between those two. Finally, the U-MD simulations were too short to establish equilibrium conformational populations, and chloroform serves only as an approximate proxy for the nonpolar interior of a lipid bilayer rather than as a model of a membrane.

## 5 Conclusion

Our findings suggest that the concept of a permeable-like conformation should be reconsidered. Compact, low-polarity conformations were accessible to every PROTAC we examined, irrespective of permeability class, so the ability to adopt such a state carries little discriminating information on its own. In the solvated simulations, what distinguished the permeable compound was not the compactness of its folded state but its persistence: it occupied a closed-flat ensemble in both water and chloroform, in two structurally distinct arrangements, whereas the moderately permeable compound did not retain its fold in either solvent, and the non-permeable compound retained one only in water. We therefore propose that chameleonicity is better understood as the persistence and controlled interconversion of a small number of well-defined conformational states than as a general capacity to become more compact, something we have found very difficult to capture with descriptors in this work. Efficient permeation may also depend on avoiding interfacial trapping and overcoming high free-energy barriers within the membrane core [63]. Moreover, some PROTACs may benefit from adopting relatively extended conformations in the membrane interior, where the energetic penalty associated with increased polar exposure may be partly offset by greater hydrophobic surface exposure [63]. Enhanced-sampling MD using the radius of gyration as collective variable, provides a workflow for generating PROTAC conformational ensembles that we expect to be useful in degrader design. Practically, maximum ring size provides a means of identifying compact states within large conformational ensembles by locating folded conformers stabilized by intramolecular hydrogen bonds. It carries no information about permeability on its own, and in this study its role was to enable the selection of starting structures with comparable compactness across all three permeability classes, reducing the influence of initial compactness on differences between the solvated ensembles. Molecular size and flexibility nevertheless remain potential confounders, reflecting an inherent limitation of the available dataset. Whether conformational persistence in bulk solvent predicts behavior within a lipid bilayer remains an open question. Addressing this question will require explicit membrane simulations, beyond the scope of the present study. Consistent with our findings, Le Manach et al. [20] showed that high molecular chameleonicity does not necessarily translate into improved oral absorption, as assessed *in vivo* using the mouse fraction absorbed (mFabs). Many poorly absorbed compounds displayed high chameleonicity according to Chamelogk, while several less-chameleonic compounds showed more favorable absorption. Compounds with comparable lipophilicity and polarity but different Chamelogk values also exhibited similarly poor absorption [20]. However, unlike PAMPA, mFabs reflects the overall intestinal absorption process, and the outcome may also be influenced by efflux, metabolism, or active protein-mediated transport. For example, conformational adaptability may affect cellular uptake through mechanisms other than passive diffusion. This distinction is illustrated by MZ1 (PROTAC-14), which is classified as chameleonic (Chamelogk = 1.5) but showed very low passive PAMPA permeability (0.1 nm/s). Nevertheless, MZ1 has been reported to partly rely on interferon-inducible transmembrane proteins (IFITMs) for cellular entry [64].

## Author contributions

**FJ:** Conceptualization, Data curation, Formal analysis, Investigation, Methodology, Software, Validation, Visualization, Writing – original draft. **RM:** Conceptualization, Funding acquisition, Resources, Supervision, Writing – review & editing.

## Acknowledgments

FJ and RM acknowledge the funding provided by the Wallenberg AI, Autonomous Systems, and Software Program (WASP), supported by the Knut and Alice Wallenberg Foundation. Computational resources were provided by the National Academic Infrastructure for Supercomputing in Sweden (NAISS), funded by the Swedish Research Council. Computational resources were provided on the Berzelius system funded by the Knut and Alice Wallenberg foundation and operated by NAISS. The authors declare no competing interests.

## Data availability

The Jupyter notebooks containing the analyses used to reproduce the reported results and figures are available at https://github.com/FJalalypour/protac-chameleonicity. Results and trajectories are archived on Zenodo at https://zenodo.org/uploads/22035338.

## Appendix

### 1. Solvent-accessible surface area (SASA) across the PROTAC permeability classes

**Figure S1.**
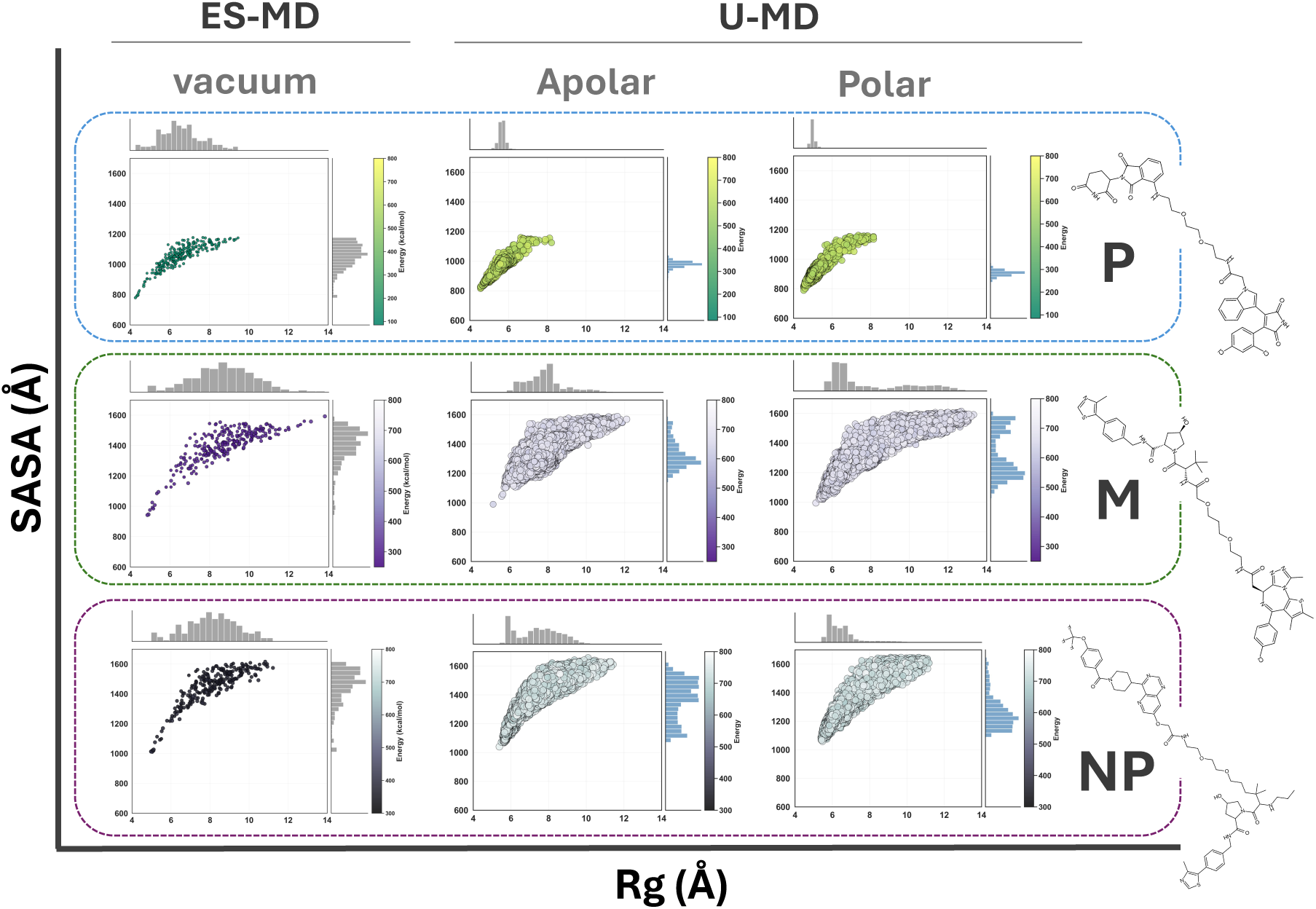
Solvent-dependent changes in solvent-accessible surface area (SASA) across PROTAC permeability classes. 2D distributions of radius of gyration (Rg) and solvent-accessible surface area (SASA) for highly permeable PROTAC-1 (P), moderately permeable PROTAC-3 (M), and non-permeable PROTAC-18 (NP). Left: conformational ensembles generated by ES-MD. Middle and right: ensembles sampled during all-atom U-MD in apolar (chloroform) and polar (water) environments, respectively. Each point represents a sampled conformation, with point color indicating its energy; energy values are interpreted within each individual PROTAC and are not directly comparable between different molecules. Marginal histograms show the corresponding distributions of Rg and SASA.

### 2. Intramolecular hydrogen-bond profile across the PROTAC dataset

**Figure S2.**
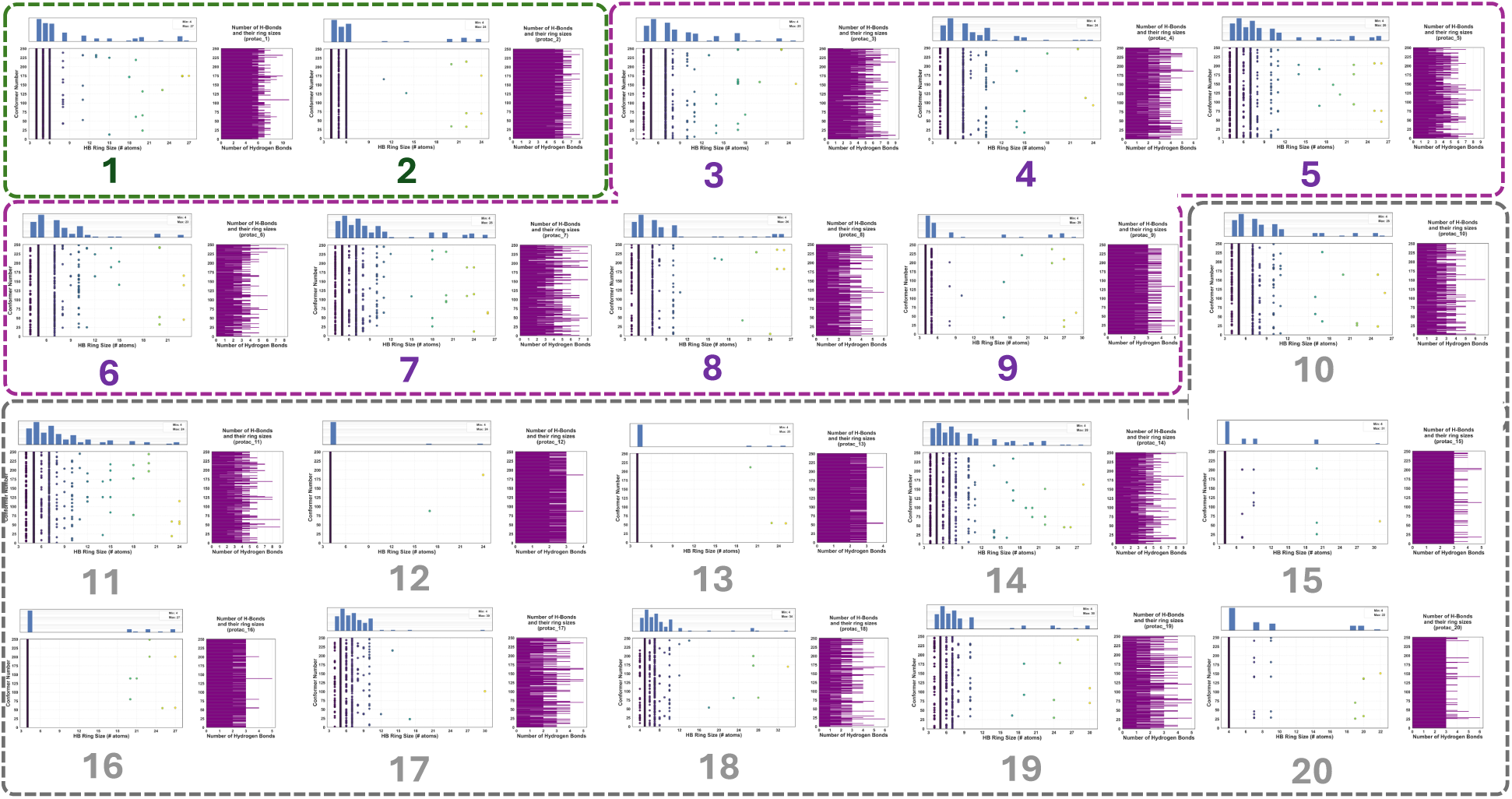
Intramolecular hydrogen-bond ring sizes across the PROTAC dataset. IHB ring sizes and the number of IHBs per conformer across ES-MD ensembles of 20 PROTACs, using a 3 Å H···acceptor distance cutoff.

**Figure S3.**
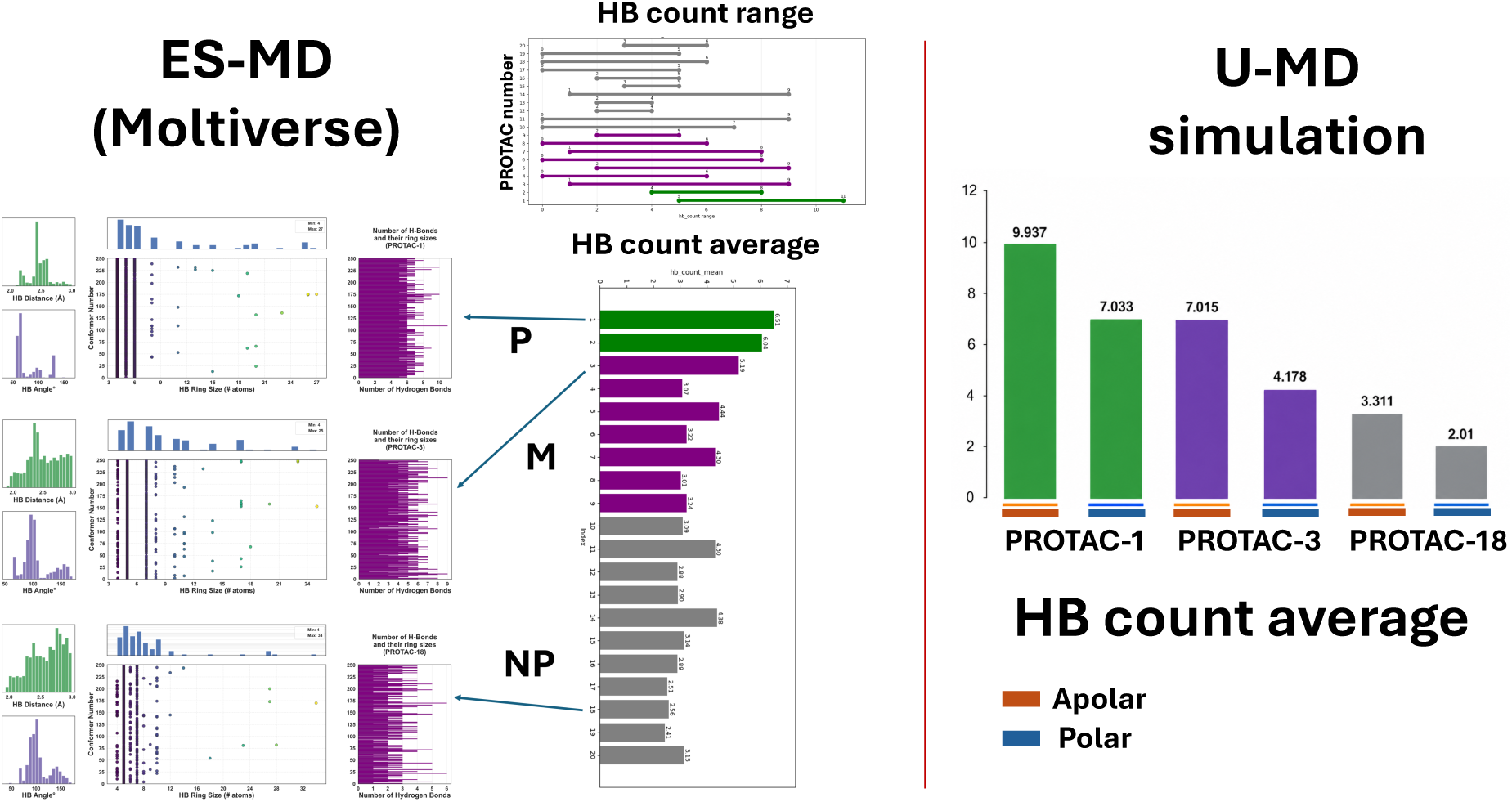
Intramolecular hydrogen-bonding across ES-MD and U-MD ensembles. IHB count ranges and average IHB counts across the 20-PROTAC ES-MD ensembles (left), with comparison of average IHB counts for PROTAC-1, PROTAC-3, and PROTAC-18 during U-MD in apolar and polar environments (right).

### 3. RMSD profiles of four independent 200ns U-MD replicas

**Figure S4.**
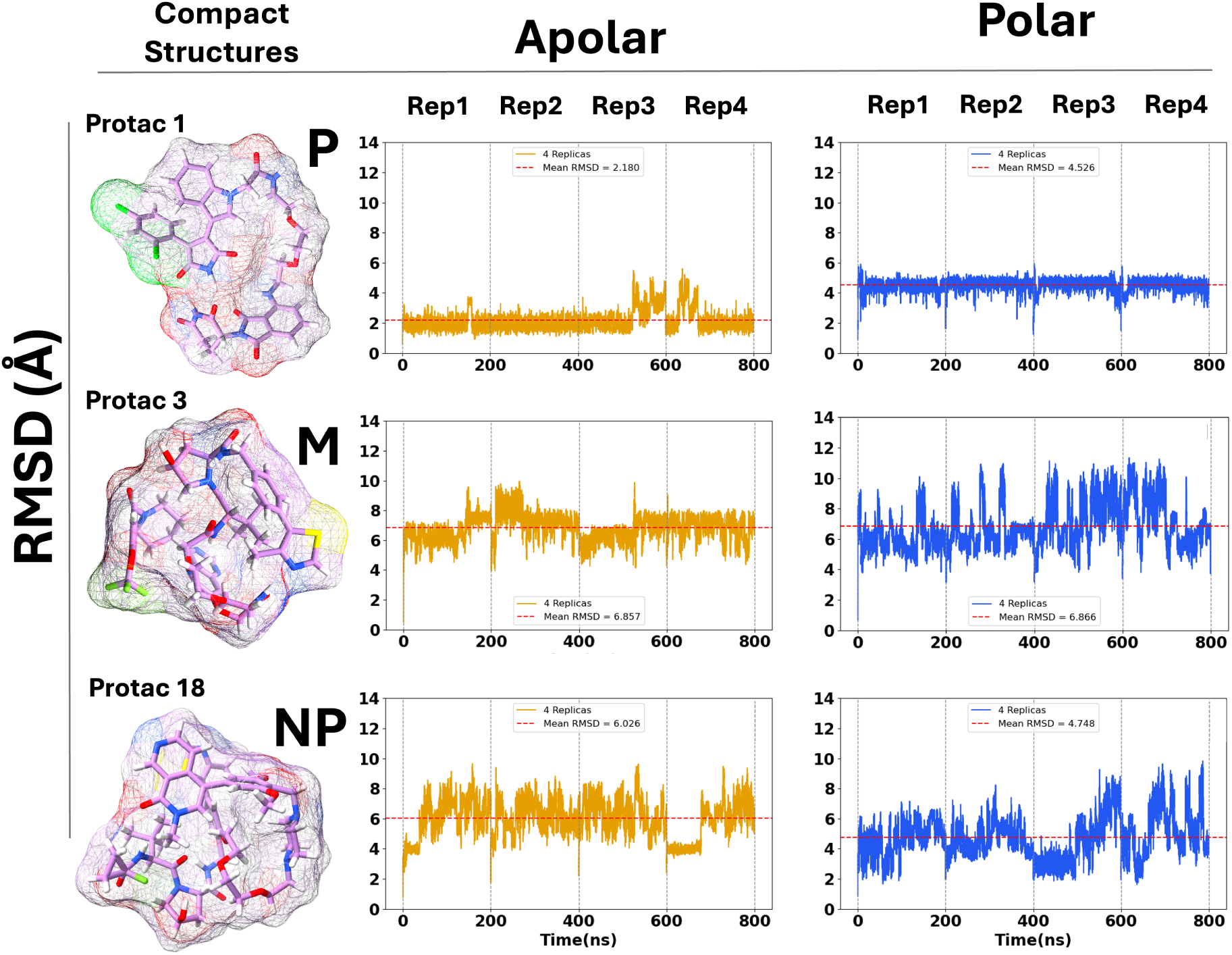
Replica-resolved conformational stability of compact PROTAC structures during U-MD. RMSD profiles of four independent 200 ns replicas for PROTAC-1 (P), PROTAC-3 (M), and PROTAC-18 (NP) in apolar (chloroform) and polar (water) environments. RMSD was calculated relative to the corresponding compact initial structure. Dashed red lines indicate the mean RMSD across the four replicas

### 4. Comparison of ES-MD (Moltiverse) and U-MD conformational ensembles and solvent-dependent U-MD distributions

**Figure S5.**
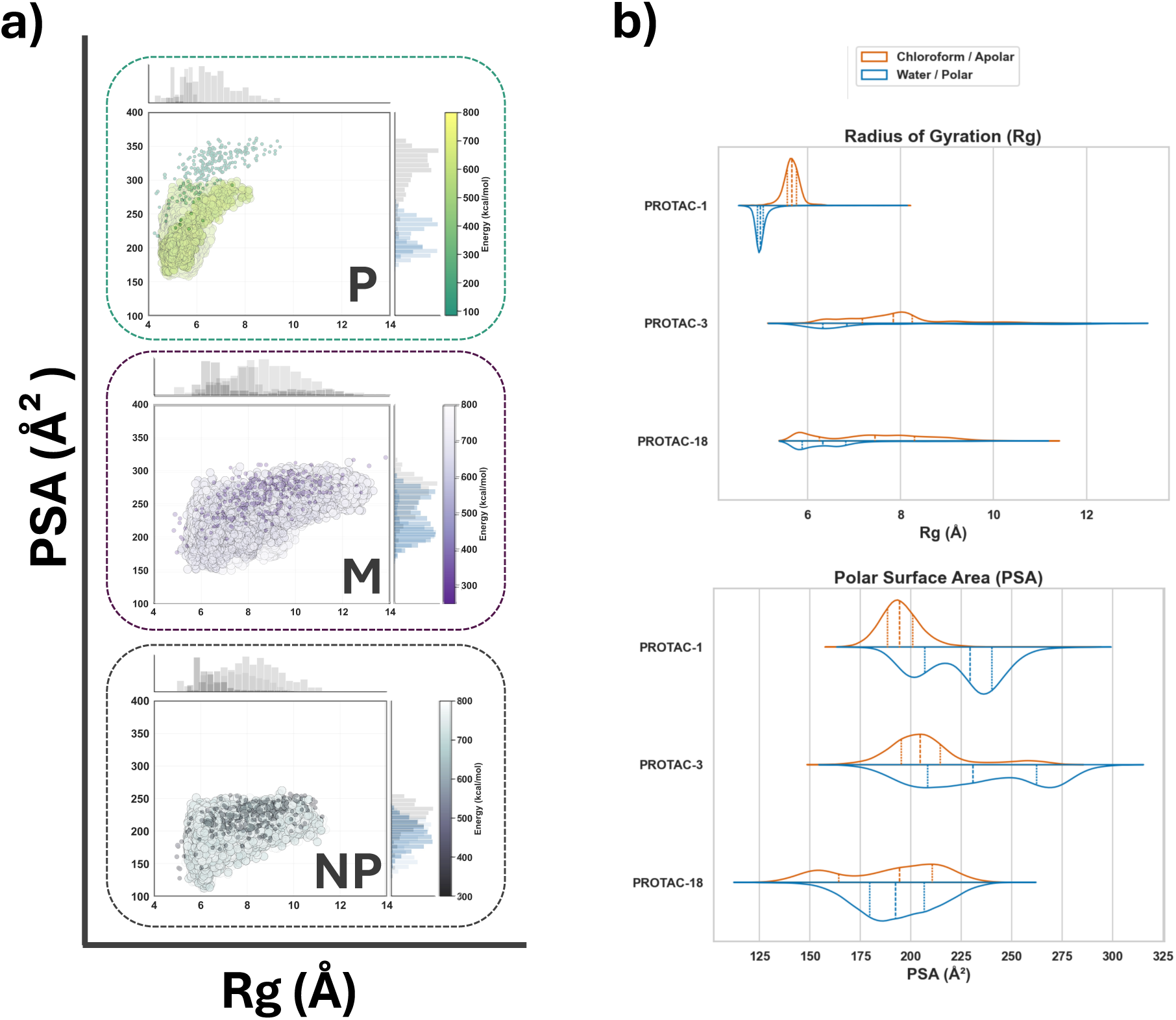
Comparison of conformational sampling by Moltiverse and U-MD. (a) Rg–PSA distributions showing partial overlap between U-MD and the broader Moltiverse ensembles. (b) Rg and PSA distributions from U-MD in apolar (chloroform) and polar (water) environments, highlighting solvent-dependent conformational sampling.

### 5. Solvent-dependent conformational shape ensembles sampled by U-MD

**Figure S6.**
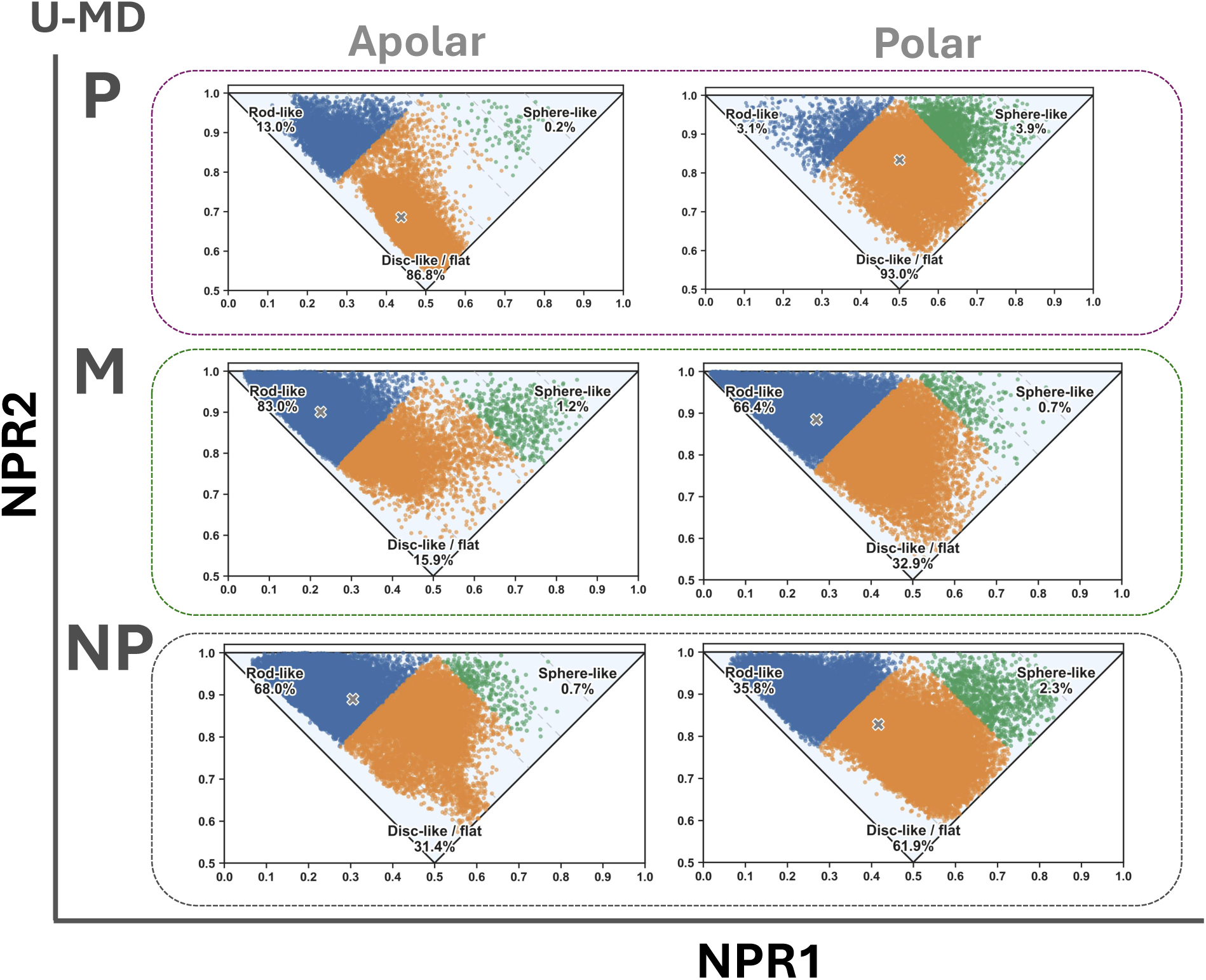
Solvent-dependent molecular shape distributions during U-MD simulations. Normalized PMI (NPR1–NPR2) distributions of PROTAC-1 (P), PROTAC-3 (M), and PROTAC-18 (NP) in apolar (chloroform) and polar (water) environments. Conformations are classified as rod-like, disc-like/flat, or sphere-like, with percentages indicating the population of each shape class.

